# Silencing of *ApoE* with Divalent siRNAs Drives Activation of Immune Clearance Pathways and Improves Amyloid Pathology in Mouse Models of Alzheimer’s Disease

**DOI:** 10.1101/2022.06.28.498012

**Authors:** Chantal M. Ferguson, Samuel Hildebrand, Bruno M.D.C Godinho, Julianna Buchwald, Dimas Echeverria, Andrew Coles, Anastasia Grigorenko, Lorenc Vanjielli, Jacquelyn Sousa, Nicholas McHugh, Matthew Hassler, Francesco Santarelli, Michael T. Heneka, Evgeny Rogaev, Anastasia Khvorova

## Abstract

The most common genetic risk factor for late-onset Alzheimer’s disease (AD) is the *APOE4* allele, with evidence for gain- and loss-of-function mechanisms. *ApoE* knockout in mice abrogates AD phenotypes but causes severe atherosclerosis due to the role of liver *ApoE* in cholesterol homeostasis. Previous attempts to inhibit brain-specific *ApoE* with anti-sense oligonucleotides only modestly reduced *ApoE* expression and had no effect on amyloid burden in adult AD mice. Here, we optimized a divalent small interfering RNA (di-siRNA) to selectively and potently silence *ApoE* in the brain. Silencing brain *ApoE* in AD mice significantly reduced amyloid plaque formation without affecting systemic cholesterol levels, confirming that brain and liver APOE pools are spatially and functionally distinct. Mechanistically, APOE appears to be a scaffold for beta-amyloid aggregation that limits clearance by microglia. Di-siRNAs from this study can be taken to pre-clinical and clinical trials to accelerate development of AD-modifying therapies and establish siRNA-based modulation of *ApoE* as a viable path towards therapeutic development.

---

Alzheimer disease (AD) is a progressive, multifactorial neurodegenerative condition for which current medications provide only short-term alleviation of symptoms (Amir Qaseem, 2008; Nhi-Ha Trinh et al., 2003). In late-onset AD, the *APOE4* allele is the most common genetic risk factor, and its presence is associated with an increase in AD incidence and a decrease in age of clinical onset (E. H. Corder, 1993; Giovanni B. Firsoni, 1995). *ApoE* is mainly produced in the central nervous system (CNS) and liver. In the CNS, glial cells (and to a lesser extent, neurons) express *ApoE* which transports lipids between cells and modulates the inflammatory response (Robert E. Pitas, 1987) (Qin Xu, 2006). Liver *ApoE* facilitates lipid uptake into peripheral tissues via low-density lipoprotein (LDL) receptors (Innerarity, 1983; Michael S. Brown, 1986). Despite decades of compelling research into the role of *ApoE* in neurodegeneration, the exact mechanism remains unclear. *ApoE* is implicated in protein clearance, immune activity, and lipid homeostasis; however, a clear-cut explanation for how *ApoE* alleles and/or levels impact AD, and consensus on whether it confers a gain or loss of function phenotype, does not exist.

Genetic removal of *ApoE* in AD models reduces beta-amyloid (Aβ) plaques – a characteristic feature of AD – and improves cognitive outcomes (David M. Holtzman, 1999; 2000; Kim et al., 2011; Liu et al., 2017; Ulrich et al., 2018). Complete loss of *ApoE* however causes spontaneous atherosclerosis in mice (Sunny H. Zhang, 1992). The tissue-specific effects of *ApoE* knockout are likely due to the differential roles of liver and CNS *ApoE*. Previous success in genetically manipulating liver-specific or brain-specific *ApoE* in mice suggests that *ApoE* pools are independent (Huynh et al., 2019; Lane-Donovan et al., 2016). However, the effects of conditional and complete *ApoE* modulation after birth—a therapeutically-relevant paradigm—are less clear.

Reduction of brain *ApoE* (∼50% of normal levels) using antisense oligonucleotides (ASO) had no significant impact on neuropathology in adult AD mice, and neonatal treatment in models of AD was necessary to produce any significant effects (Huynh et al., 2017). Although this could suggest that embryonic *ApoE* silencing is required to reduce AD phenotypes, it is also possible that residual (∼50%) ApoE expression in the CNS or from the systemic circulation suffices to maintain pathology. Similarly, removing one copy of *ApoE* in AD mice only modestly improves amyloid pathology (Bales et al., 1997). Conversely, in models of tauopathy, 50% reduction in ApoE was sufficient to reduced markers of neurodegeneration (Litvinchuk et al., 2021). Thus, ameliorating AD pathology might require near complete silencing of *ApoE* in adult brain.

Small interfering RNAs (siRNAs) guide potent and sustained gene silencing in vivo via an RNA-induced silencing complex (RISC), and the chemical architecture of an siRNA determines where in the body it will be delivered (Khvorova and Watts, 2017). For example, GalNAc-conjugated siRNAs selectively deliver to liver hepatocytes (Akinc et al., 2010; Nair et al., 2017; Nair et al., 2014), where a single injection supports 6–12 months of clinical efficacy (Ray et al., 2017). We have recently described a divalent (di)-siRNA scaffold that is efficiently taken up by neurons and glial cells and supports potent (>95%) and sustained (up to 6 months) silencing throughout the CNS (Alterman et al., 2019).

Here, we engineered siRNAs targeting mouse *ApoE* and achieved selective silencing of liver and CNS *ApoE* pools in APP/PSEN1 and 5xFAD mouse models. Near-complete silencing of liver *ApoE* in AD mice increased serum cholesterol with no detectable impact on brain *ApoE* or Aβ pathology. Potent reduction of CNS *ApoE*, either before or after onset of AD pathology, decreased Aβ burden with no detectable effect on systemic cholesterol. Mechanistically, *ApoE* serves as core for amyloid plaque seeding and its removal activated immune response pathways that promote Aβ clearance. Collectively, these results identify siRNA-mediated silencing of *ApoE* as a potential therapeutic for AD in a landscape where disease-modifying therapies are limited.

## Results

### CNS and systemic *ApoE* are spatially and functionally distinct in APP/PSEN1 mice

Tissue-specific *ApoE* knockouts have revealed that CNS and systemic ApoE are independently maintained in mice (Huynh *et al*., 2019; Lane-Donovan *et al*., 2016). To determine if siRNA-mediated silencing of CNS *ApoE* affects ApoE levels in AD mice without affecting systemic ApoE (expressed in liver) and vice versa, we engineered fully chemically-modified siRNAs targeting mouse *ApoE* in either liver-active (GalNAc^APOE^) or CNS-active (di-siRNA^APOE^) chemical configurations (**Extended Data Table 1**). Two-month-old APP/PSEN1 mice (JAX#34832) (Garcia-Alloza et al., 2006; Jankowsky et al., 2004; Jankowsky JL, 2001) were injected intracerebroventricularly (ICV) with 237 μg of di-siRNA^APOE^ or non-targeting control di-siRNA^NTC^. A second group was injected subcutaneously (SC) with 10 mg/kg GalNAc^APOE^ or GalNAc^NTC^. Two-months post-injection we measured *ApoE* mRNA and protein and serum cholesterol levels in brain and liver (4-months old; **Fig. 1a-b, Extended Data Fig. 1a-b**). In di-siRNA^APOE^-treated mice, we observed >90% *ApoE* mRNA and >95% protein silencing (p<0.0001) in the brain and ∼50% silencing in liver (p<0.0001) compared to di-siRNA^NTC^-treated mice (**Fig. 1c-d**). In GalNAc^APOE^-treated mice, we observed >99% ApoE mRNA and protein silencing in the liver (p<0.0001) and no silencing in brain (**Extended Data Fig. 1c-d, Extended data Fig. 2 for raw western blots**). Despite the partial reduction of liver *ApoE* in di-siRNA^APOE^ -treated mice, serum cholesterol levels remained normal **(Fig.1e)**. By contrast, the near-complete silencing of liver *ApoE* in GalNAc^APOE^-treated mice was associated with increased LDL-cholesterol levels compared to control GalNAc^NTC^-treated mice (∼300%, p<0.001) (**Extended Data Fig. 1e**). We observed no liver toxicity di-siRNA^APOE^-treated or GalNAc^APOE^-treated mice compared to NTC-treated mice **(Extended Data Table 2).** These data indicate that, at least in the short-term, CNS ApoE does not rescue the loss of systemic ApoE or regulate systemic cholesterol levels, and systemic ApoE does not rescue the loss of brain ApoE. The ability to silence CNS *ApoE* without affecting the systemic ApoE or cholesterol levels allows us to explore whether siRNA-mediated silencing of CNS *ApoE* affects AD pathology.

**Figure 1:**
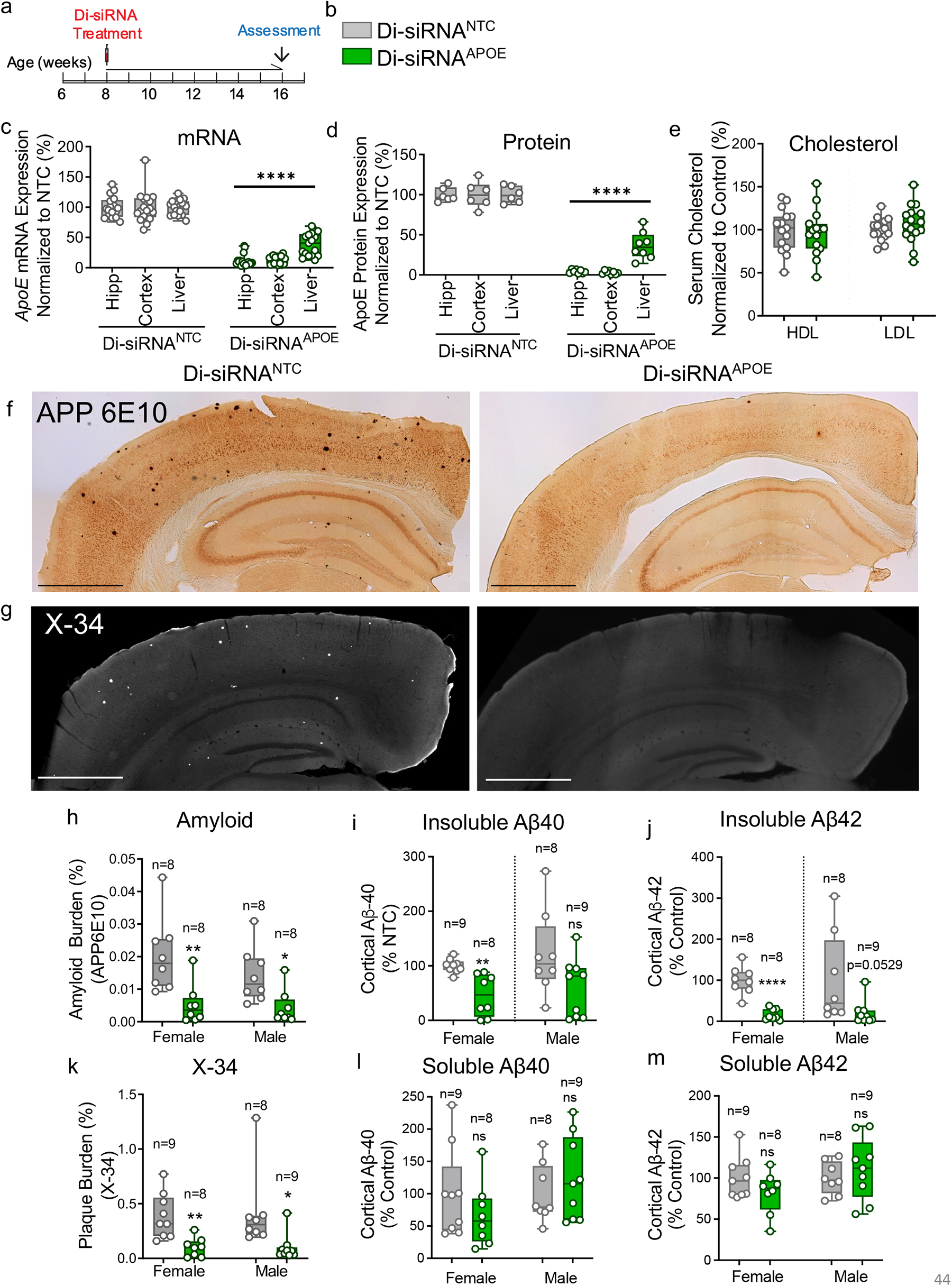
Silencing brain *ApoE* reduces AD neuropathology in APP/PSEN1 mice with no effect on serum cholesterol. (**a, b**) Experimental timeline. di-siRNA dose: 237 μg. Di-siRNA^NTC^ shown in grey, and di-siRNA^APOE^ shown in green. *ApoE* mRNA (**c**) and (**d**) protein expression in the hippocampus, cortex, and liver 2-months post administration of di-siRNA^NTC^ or di-siRNA^APOE^. (**e**) Serum HDL and LDL cholesterol in di-siRNA^NTC^ and di-siRNA^APOE^ mice (n=8-9 per group). (**f**) APP6E10 plaques in the cortex 2-months post injection of in treated and controls. (**g)** X-34-positive plaques in cortex 2-months post treatment. (**h**) Sex-stratified quantification of APP6E10-positive cortex plaques. (**i**) Water-insoluble Aβ-40 fibrils in cortex samples. (**j**) Water-insoluble Aβ-42 fibrils in cortex samples. (**k**) Sex-stratified quantification of X-34-positive plaques in cortex. (**l**) Soluble Aβ-40 fibrils in cortex samples. (**m**) Soluble Aβ-42 fibrils in cortex samples. mRNA evaluated using QuantiGene. APOE protein quantified with WES ProteinSimple; Statistical analysis: one-way ANOVA or T-test using GraphPad Prism. Error bars are SD. (****p <0.0001; ***p<0.002; **p<0.01; *p<0.05).

### CNS-selective silencing of *ApoE* reduces amyloid pathology in APP/PSEN1 model

To evaluate the effects of *ApoE* silencing on AD pathology in APP/PSEN1 mice, we measured the number and size of amyloid plaques at two-months post-injection. Toxic amyloid fragments (Aβ-40, Aβ-42) aggregate into plaques that are visible throughout the brain on post-mortem histology. Amyloid plaques were detected by immunohistochemistry (IHC) for amyloid (APP6E10) or by X-34 staining (which is highly sensitivity for β-pleated amyloid) (Ikonomovic et al., 2006), and amyloid plaque burden was calculated as percent area positive for amyloid plaques (see Methods for details). Compared to di-siRNA^NTC^, di-siRNA^APOE^ reduced amyloid plaque burden throughout the brain analyzed by APP6E10 immunohistochemistry (female: 86%, p<0.0001; male: 70%, p=0.0158) (Fig. 1f, h) and X-34 staining (female: 78%, p<0.0001; male: 79%, p=0.0029) (Fig. 1g, k). In addition to visible aggregated plaques, amyloid burden can be measured using quantitative ELISA based assays specific for Aβ-42 or Aβ-40 fragments (Gu and Guo, 2013; Han et al., 2017). We used ELISA to measure Aβ levels in soluble and insoluble fractions of cortical lysates. Consistent with the reduced amyloid plaque burden observed in staining studies, silencing *ApoE* resulted in reduced levels of insoluble Aβ-40 (female: 56%, p=0.011; male: 40%, p=0.0738) (**Fig. 1i**) and Aβ-42 (female: 84%, p<0.0001; male: 80%, p=0.0529) (**Fig. 1j**). Soluble Aβ-42 and Aβ-40 levels were similar between treatment groups (**Fig. 1l, m**). Consistent with previous data (Ordonez-Gutierrez et al., 2016), the degree of amyloid burden differed between female and male mice, but di-siRNA^APOE^ treatment reduced amyloid burden pathology similarly in both sexes. Importantly, near-complete silencing of liver *ApoE* had no effect on amyloid burden or insoluble Aβ-42 levels but did modestly increase soluble Aβ-42 (**Extended Fig. 1e-i**). Collectively, our results show that near-complete silencing of CNS *ApoE* (but not liver *ApoE*) before the onset of amyloid pathology significantly reduces amyloid burden in adult APP/PSEN1 mice without affecting systemic cholesterol homeostasis.

### Silencing *ApoE* After Onset of Pathological Changes Reduces Amyloid Pathology in 5xFAD mice

To study the effects of silencing *ApoE* after the onset of amyloid pathology, we turned to the 5xFAD mouse model, which expresses humanized *APP* harboring 3 mutant alleles—Swedish (K670N, M671L), Florida (I716V), and London (V717I) alleles—and human *PSEN1* harboring two mutations (M146L and L286V). The 5xFAD mice show aggressive amyloid deposition, astrocyte activation, and neuronal cell death beginning between 6 and 8 weeks of age (Oakley et al., 2006). We injected (ICV) 9-week-old 5xFAD mice with 237 μg of di-siRNA^APOE^ or di-siRNA^NTC^ (control) and measured brain *ApoE* expression, amyloid burden, and transcriptome changes at two weeks and two months post-injection (**Fig. 2a-b**). Di-siRNA^APOE^ potently silenced *ApoE* mRNA in all regions of the brain (**Fig. 2c**) (>95%, p<0.0001) at both timepoints. At these early time points, 5xFAD mice do not exhibit significant AD-related cognitive deficits. However, potent silencing of *ApoE* did not impair cognitive function in open field, novel object recognition (NOR), or y-maze spontaneous alternation tests at two months post-injection (**Extended Data Fig. 3**). While we were unable to determine the impact on AD-related cognitive changes from these experiments, these results indicate that silencing CNS *ApoE* with siRNAs is well-tolerated in adult AD mice. Further studies are needed in older mice to evaluate the impact of silencing *ApoE* on the cognitive effects of AD.

**Figure 2:**
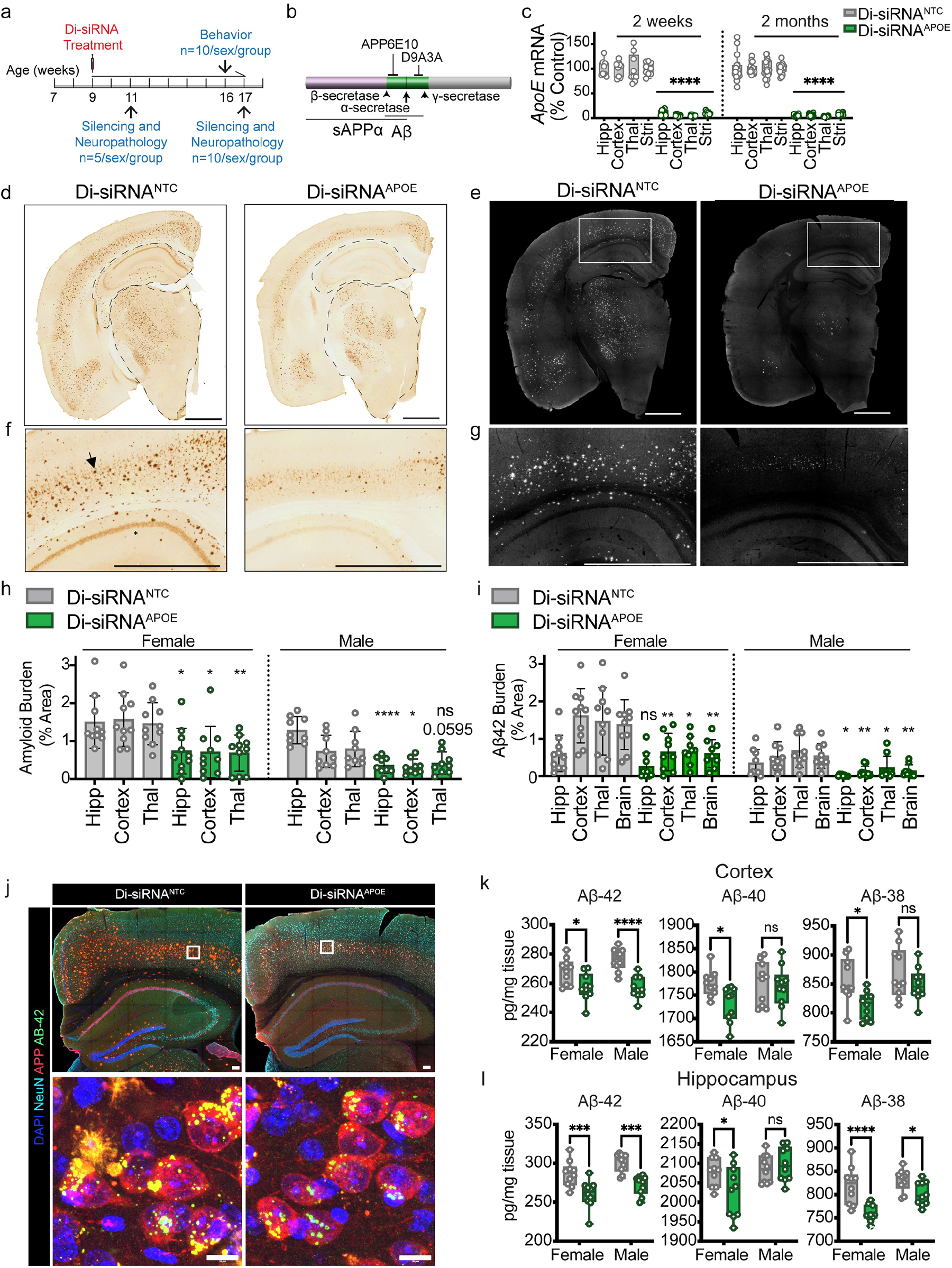
Silencing *ApoE* after the onset of pathologic changes improves amyloid burden in 5xFAD mice. (**a**) Experimental design. (**b**) Schematic showing antibody binding sites for APP6E10 and Aβ-42-specific antibody (D9A3A). APP6E10 will react with Aβ and soluble APPα (sAPP α). (**c**) mRNA expression throughout the brain 2-weeks and 2-months post administration of di-siRNA^APOE^. Controls (di-siRNA^NTC)^ shown in grey and treated (di-siRNA^APOE)^ shown in green. (**d**) Amyloid deposition using APP6E10 in controls (left) and treated (right) animals. Scale bar: 1000 μm. (**e**) Aβ-42 using D9A3A in controls (left) and treated (right) animals. (**f**) Zoom of cortical regions from (d) showing neuronal amyloid reactivity in both groups and reduction in amyloid plaque deposition in di-siRNA^APOE^ treated animals. (**g**) Zoom of cortical regions from (e) showing Aβ-42 plaque reduction in di-siRNA^APOE^ treated animals. Scale bar: 1000 μm (**h-i**) Quantification of plaque burden measured as percent area, separated by sex. (**j**) co-labeling of amyloid (red, APP6E10), Aβ-42 (green), neurons (light blue, NeuN), and cell nuclei (dark blue, Dapi) in controls (left) and treated (right) 17-week old 5xFAD mice. Top: 10x resolution, scale bar: 100 μm. Bottom: High-resolution imaging showing presence of intraneuronal Aβ-42 and amyloid; 40x, scale bar: 10 μm. (**k**) MDS Quantification of insoluble amyloid in the cortex and (**l**) hippocampus. Statistics: t-test per brain region; sex separated. *: p<0.0332; ** p<0.0021; *** p<0.0002; **** p<0.0001. n=9-10 per sex per group. Timepoint: 17 weeks old, 2 months post injection.

At two weeks post-injection, both di-siRNA^APOE^- and di-siRNA^NTC^-treated animals showed early signs of amyloid pathology, with the most extensive plaque deposition in the posterior hippocampus and entorhinal cortex, most reactivity associated with intraneuronal APP, and extracellular plaque deposition primarily in the CA1 hippocampal region (**Extended Data Fig. 4a-c**).

At two months post-injection, compared to controls, we observed significantly less amyloid burden in the brains of di-siRNA^APOE^-treated mice (**Fig. 2d, f, h**) (**Extended Data Table 3**). In di-siRNA^APOE^-treated male but not female mice, amyloid plaques in the hippocampus and cortex were smaller than those in control mice (p=0.0034 and p=0.0326, respectively) (**Extended Data Fig. 5a**). Differences in plaque size were not observed in other regions of the brain where fewer plaques were detected. The number of amyloid plaques was significantly reduced in all brain regions in female mice (hippocampus: p=0.0023; cortex: p=0.0019; thalamus: p=0.0004), and in the thalamus of male mice (p=0.0286) (**Extended Data Fig. 5b, Extended Data Table 3**).

The APP6E10 antibody used to measure amyloid burden detects Aβ fragments (Aβ 38, 40, 42) and the normally processed 100-kDa APP product (**Fig. 2b**) (Youmans et al., 2012). Indeed, IHC using APP6E10 strongly stained extracellular plaques (black arrow) and diffuse intraneuronal APP in cortical and hippocampal neurons (black Asterix) (**Fig. 2f**). To differentiate between general amyloid burden measured by APP6E10 and Aβ-specific plaque load, we performed immunofluorescence with an Aβ-42 specific antibody (Cell signaling; D9A3A). *ApoE* silencing led to a lower amyloid plaque burden (**Fig. 2e, g, i**) and fewer amyloid plaques (**Extended data Fig. 5c**) throughout the brain. Plaque size however was unaffected by *ApoE* silencing (**Extended data Fig. 5d, Extended Data Table 4)**.

Using multiplex-immunofluorescence to visualize Aβ-42 and APP in 5xFAD mice, we observed three amyloid-related molecular signatures: 1) neuron-specific diffuse intracellular 100-kDa APP staining (**Fig. 2j, red staining**); 2) large extracellular Aβ-42 and APP-positive plaques (yellow); and 3) neuron-specific intravesicular Aβ-42 foci (green). Di-siRNA*^APOE^* treatment had no obvious effect on diffuse intracellular 100kD APP (**Fig. 2j, red**), partially reduced intraneuronal perinuclear Aβ-42 vesicular foci (**Fig. 2j, green**), and as shown above, dramatically reduced extracellular plaques by ∼80% (Fig 2). Multiplex ELISA MSD assays (MSD K15200E) revealed that whereas *ApoE* silencing had no effect on soluble Aβ-42 or Aβ-40 levels (**Extended data Figure 5e**), *ApoE* silencing reduced the burden of insoluble Aβ-42 (**Fig. 2k, l**), the more toxic and aggregate-prone fragment.

### Silencing *ApoE* reduces amyloid carboxyl-terminal fragments

In AD patients, toxic amyloid plaques accumulate when the production of amyloid protein fragments is not balanced by clearance (Kummer and Heneka, 2014). Amyloid precursor protein processing on the cell membrane produces carboxy-terminal fragments (α-CTFs) by a non-amyloidogenic pathway, and β-CTFs via an amyloidogenic pathway (Kummer and Heneka, 2014). In healthy brains, amyloid fragments are cleared by amyloid degrading enzymes (ADEs), including neprilysin, insulin-degrading enzyme (IDE), and matrix metalloproteinases (Leissring et al., 2003; Miners et al., 2011). To determine how silencing *ApoE* helps improve amyloid pathology, we measured α- and β-CTFs in hippocampal and cortical brains samples (Haass et al., 2012). In control animals, α- and β-CTF levels increased between timepoints, consistent with worsening pathology as animals age (**Fig. 3a-d**). After 2 months of silencing CNS *ApoE*, however, cortical and hippocampal Aβ-42 was markedly reduced (**Fig. 3a-d**), and α- and β-CTFs were markedly reduced in the cortex, suggesting that silencing *ApoE* reduces the production of amyloid fragments via both amyloidogenic and non-amyloidogenic pathways. Silencing *ApoE* did not significantly affect mRNA levels of APP, presenilin (PSEN1/PSEN2), or the APP cleaving enzyme BACE1 (**Fig. 3e**). These data suggest that silencing CNS *ApoE* does not reduce amyloid burden by disrupting the expression of key APP processing machinery.

**Figure 3:**
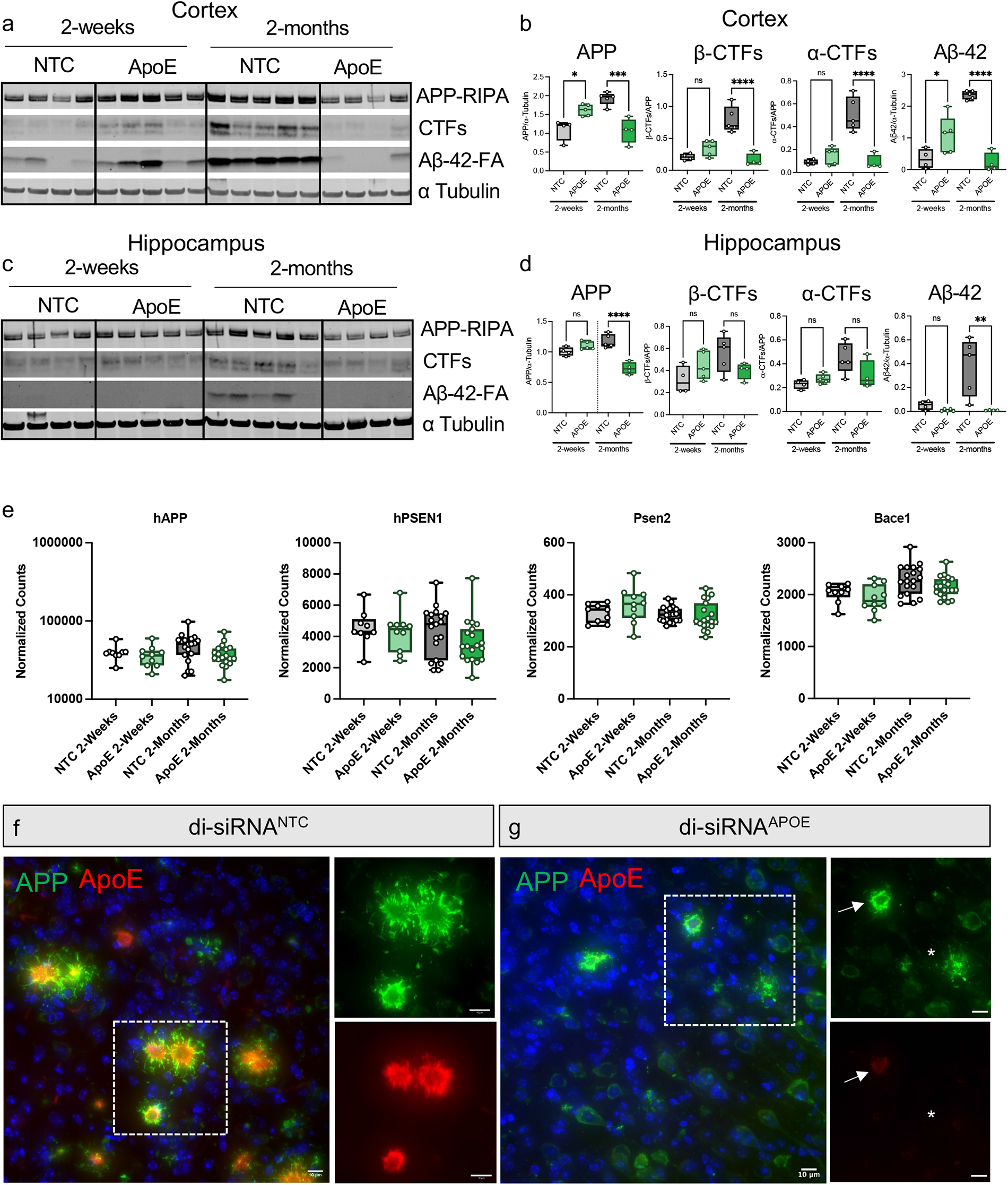
Silencing ApoE decreases amyloid CTF production and removes structural support for amyloid plaques. (a) Western blots and quantification (**b**) showing reduction in CTFs and Aβ-42 in the cortex of treated compared to control animals. **(c)** Western blots and quantification (**d**) showing reduction in APP and Aβ-42 in the hippocampus of treated compared to control animals. N=4-5 per group. Statistics: one-way ANOVA: p<0.0332; ** p<0.0021; *** p<0.0002; **** p<0.0001. **(e)** RNA expression levels of APP, PSEN1, PSEN2, and BACE1 in the cortex of treated and control animals. (**f-g**) Co-staining of amyloid plaques (APP6E10, green) and ApoE (red) in control (f) and *ApoE* silenced (g) 5xFAD mice. Insets show amyloid (top, green) and ApoE (red, bottom). Arrows denote early plaque with ApoE present in core while star denotes ApoE poor plaque with altered structure. 63x, scale bar: 10 μm.

### *ApoE* core defines amyloid plaque structure and limits access to glia-mediated clearance

We next wondered if silencing *ApoE* reduces amyloid plaques by preventing ApoE from promoting aggregation of amyloid fragments. Previous studies have observed that ApoE is enriched in the core of amyloid plaques, suggesting that ApoE serves as a scaffold for the aggregation of amyloid fragments into non-degradable plaques and that loss of the ApoE core allows phagocytic clearance of plaques by microglia and turnover of toxic amyloid fragments (Hashimoto et al., 2012; Huynh *et al*., 2017; Kiskis et al., 2015; Meilandt et al., 2020; Parhizkar et al., 2019; Sala Frigerio et al., 2019; Stephen et al., 2019; Ulrich *et al*., 2018). Indeed, we also observed large, ApoE-rich amyloid plaques in controls and a reduction in ApoE-rich plaques after the therapeutic silencing of *ApoE* (**Fig. 3f-g**). The ability to reverse plaque formation by silencing *ApoE* strongly suggests that ApoE serves as a scaffold for amyloid plaque formation. We hypothesize, as others have previously, that silencing *ApoE* destroys the core, allowing phagocytic clearance of plaques by microglia and the turnover of toxic amyloid fragments (Fitz et al., 2020; Sala Frigerio *et al*., 2019; Stephen *et al*., 2019; Ulrich *et al*., 2018). We build upon these studies by showing that altering the ApoE rich core and thus amyloid plaque formation can be achieved with therapeutic silencing of *ApoE*.

In early stages of AD, amyloid degrading enzymes released by microglia cannot keep pace with the production and aggregation of toxic amyloid fragments (Rapic et al., 2013). In control di-siRNA^NTC^-treated mice, IBA1-positive microglia clustered around plaques (**Fig. 4a-b**). In di-siRNA^APOE^-treated mice, however, microglial were present but formed fewer clusters around amyloid plaques (**Fig. 4c**). Microglia appeared to form fewer branches or processes in control animals, suggesting that *ApoE* silencing increases ramification. However, the dense clustering of microglia and complexity of the tissue samples made it difficult to quantify. The effect of *ApoE* silencing is likely to be multifaceted: making plaques more accessible to microglia causes innate immune activation, while lowering amyloid burden may reduce secondary inflammation and encourage prolonged cell survival.

**Figure 4:**
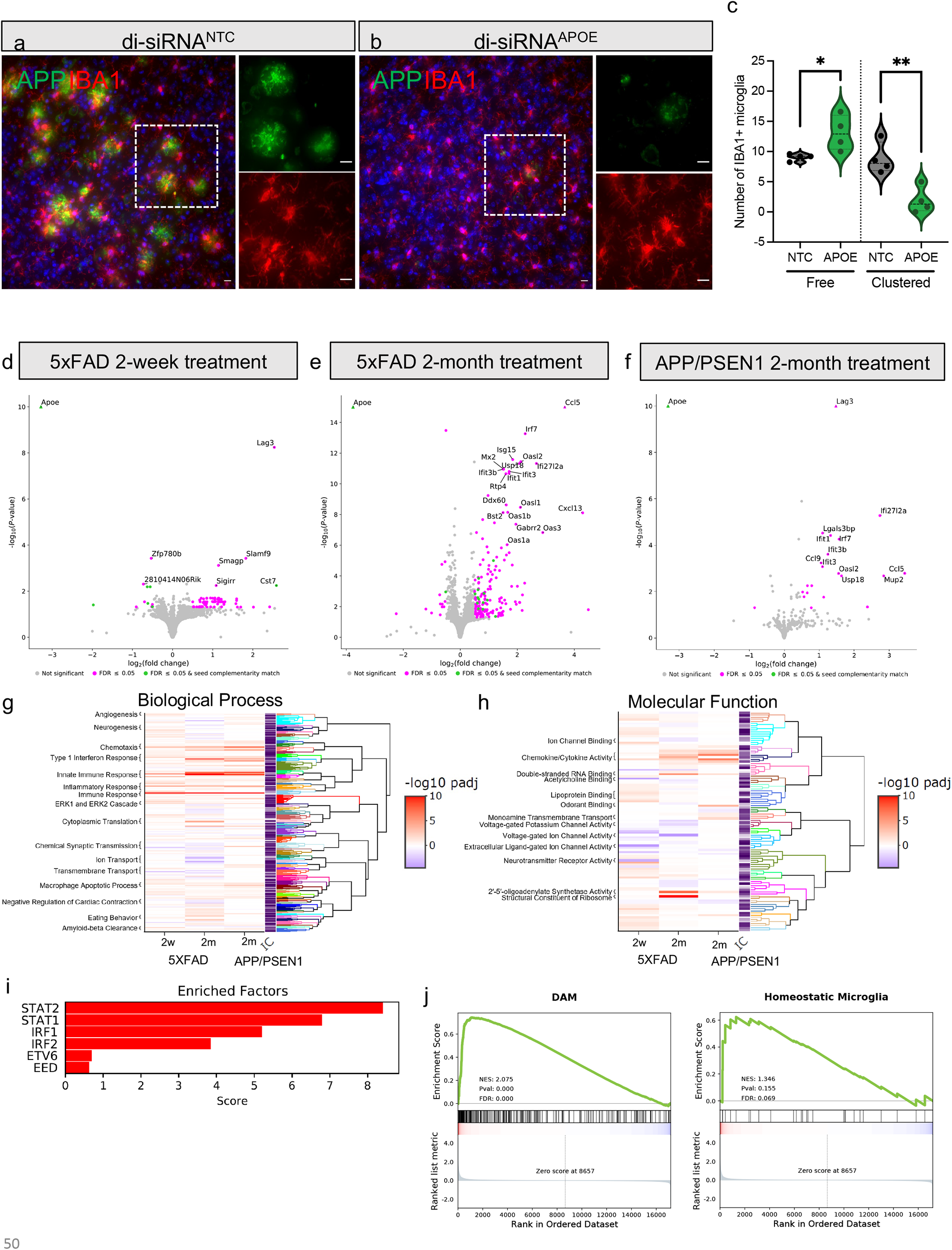
*ApoE* silencing activates expression of immune-related pathways. **(a-b)** Co-staining of amyloid plaques (APP, green) and microglia (IBA1, green) in control (**a**) and treated (**b**) 5xFAD mice. Insets show amyloid (top, green) and IBA1 (red, bottom). (**c**) Quantification of IBA1+ cell clustering around amyloid plaques. 40x, scale bar: 10 μm. Representative images from corresponding cortex regions. Statistics: One Way ANOVA. **(d-f)** Volcano plots showing differentially expressed genes after two weeks (**d**) and two months (**e**) of *ApoE* silencing in 5xFAD mice. (**f**) Differentially expressed genes after two months of *ApoE* silencing in APP/PSEN1 mice. FDR set to <0.05; shown in pink. Green denotes FDR<0.05 and presence of the seed match in the 3’UTR. Grey: not-significant. Log2fold change cut-off set to 0.5 (+/-). (**g-h**) Gene set enrichment analysis on genes expressed in all conditions, clustered by gene set member semantic similarity. Labels indicate cluster themes. (**i**) Analysis of transcription factor target gene enrichment in upregulated genes at 2 months post-treatment in 5XFAD, using the MAGIC package. (**h**) Comparison of gene overlap between *ApoE* silencing for 2-months in 5XFAD and marker genes of disease associated microglia (DAM) and homeostatic microglia from Keren-Shaul et al., 2017. 5xFAD mouse model; Age: 11 and 17 weeks old. Timepoints: 2-weeks post treatment, 2-months post injection. N=9-10 per group at 2-weeks post injection; n=18-20 per group 2-months post injection. APP/PSEN1 mouse model; Age: 16 weeks old, 2-months post treatment. N=8-10 per group.

### *ApoE* silencing activates innate immune responses

To probe the potential impact of *ApoE* silencing on immune-driven clearance, we examined the effects of *ApoE* silencing on gene expression in both APP/PSEN1 (cortex, 2 months post-injection, n=8-10/group) and 5xFAD (cortex, 2 weeks and 2 months post-injection, n=18-20/group) mouse models. *ApoE* was the most significantly downregulated gene in 5xFAD mice at 2-weeks (log2 fold-change -3.28, p=7.28e^-62^) and 2-months (log2 fold-change -3.76, p=6.63e^-165^) post-injection and in APP/PSEN1 mice (-2.85, p=2.43e^-18^) at 2-months post-injection (**Fig. 4d-f, Extended Data Table 5**). Only 2 to 4 genes with complementarity to the di-siRNA seed sequence were marginally down-regulated, perhaps related to off-target siRNA activity. The high specificity of di-siRNAs observed here is consistent with previous reports of similar type of compounds in CNS (Alterman *et al*., 2019). Thus, off-target events are unlikely to contribute to transcriptome changes.

Although we did not observe phenotypic changes in 5xFAD mice at 2-weeks post-injection, silencing *ApoE* resulted in detectable transcriptomic changes, suggesting initiation of underlying biological processes that may affect disease progression (**Fig. 4d**). After two months, transcriptome changes were more profound in both models (**Fig. 4d-f**), consistent with the significant reduction in plaque number. In both mouse models, the transcriptome effects of *ApoE* silencing were skewed towards gene activation (**Fig. 4d-f**). In 5xFAD mice, 64 genes were upregulated (log2 fold change cutoff of 0.5 and adjusted p-value cutoff of 0.05) at two weeks and 160 genes were upregulated at two months post-treatment. Conversely, only 12 and 14 genes were downregulated at the two weeks and two months. In di-siRNA^APOE^-treated APP/PSEN1 mice, 21 genes were significantly upregulated, while only 1 was downregulated. Interestingly, the transcriptional effects were proportional to the degree of pathologic improvement in treated animals compared to controls. This is consistent with studies where genetic *ApoE* knockout had much smaller effects on transcriptome signatures in healthy animals compared to animals with significant disease (Pandey et al., 2019).

Sex affects *ApoE*-related progression of AD. Nevertheless, transcriptional changes in response to *ApoE* silencing were similar in male and female mice, with ribosomal subunit protein genes being the exception (**Extended Data Fig. 8d**). Interferon genes were activated in both sexes, but the effect was stronger in male mice at 2-months post-injection (**Extended Data Fig. 8d**).

Gene set enrichment analysis (GSEA) showed a consistent increase in innate immune processes following di-siRNA^APOE^ treatment, with genes involved in type 1 interferon response and cellular response to viral pathogens being upregulated at both timepoints and in both models (**Fig. 4g-h**). At two-months post-treatment, innate immune response was the most pronounced signature, especially in 5xFAD mice. Upregulated genes included *Stat1* and *Stat2* and genes regulated by *Stat1* and *Stat2*, including oligo-adenylate synthase genes (*Oas1a, Oas1b, Oas1g, Oas2, Oas3, Oasl1, Oasl2*), interferon-related genes (*Irf7, Irf9, Stat1, Ifit1, Ifit2, Ifit3, Ifi27, Ifi35, Ifi44)*, and cytokines/chemokines (*Ccl12, Ccl2, Ccl5, Cxcl10, Cxcl13, Cxcl16, Tnfsf13b*) (**Fig. 4i, Extended data figure 7, Extended data table 5-7**) (Platanias, 2005). *ApoE* knockout induces more pronounced changes in inflammatory gene signatures in affected AD mice than in the absence of disease (i.e., wild-type animals) (Pandey *et al*., 2019), suggesting that *ApoE* silencing is not the only factor driving inflammation. Rather, the strong interferon signature associated with *ApoE* silencing might reflect underlying disease-associated microglia (DAM) or activated astrocytes (Sala Frigerio *et al*., 2019). Indeed, we observed a strong correlation with DAM expression signatures in AD mouse models identified in single-cell RNA sequencing studies (Fitz et al., 2021; Hasel et al., 2021; Mathys et al., 2017; Sala Frigerio *et al*., 2019) (**Fig. 4j, Extended Data Fig. 9**). Interestingly, *ApoE* silencing expression profiles were positively associated with the microglial response to injecting Aβ/ApoE3 lipoprotein complex into wild-type mouse brain and negatively associated with the response to injecting Aβ/ApoE4 lipoprotein (**Extended Data Fig. 9d**) (Fitz *et al*., 2021),

Pathways associated with axonogenesis, neurotransmitter transport, and synaptic organization were downregulated in 5xFAD mice after 2 months *ApoE* silencing (**Fig. 5**). The absence of this signature in APP/PSEN1 mice suggests that it could be a direct consequence of exaggerated glial response in 5xFAD mice. Two-weeks post treatment in 5xFAD mice, AD-associated genes, *Trem2*, *Slamf9, Tmem119,* and *Cst7* were upregulated. *Tmem119* remained slightly upregulated at 2-months, but just below the set threshold, while there were no significant expression differences in the other genes (**Extended data table 5**). Signatures and components of the classical complement cascade (*C1qa, C1qb, C1qc*) were upregulated, consistent with activation of synapse pruning and lysosomal activity in homeostatic microglia (Sala Frigerio *et al*., 2019), which may serve to strengthen healthy neuronal cells. Interestingly, after 2-months of *ApoE* silencing in 5xFAD mice, we observed an increased expression of other lipoproteins, including ApoA1 (log2fold change=0.79, p=0.0178) and ApoC1 (log2fold change=0.86, p=0.003), which could reflect a compensatory response to ensure proper lipid transport and cellular growth in the brain (Lefterov et al., 2010; Lewis et al., 2010). Taken together, ApoE seems to play a key role in immune system processing and silencing *ApoE* with therapeutic siRNAs may be an effective way to stimulate clearance of toxic pathogen-like proteins.

**Figure 5:**
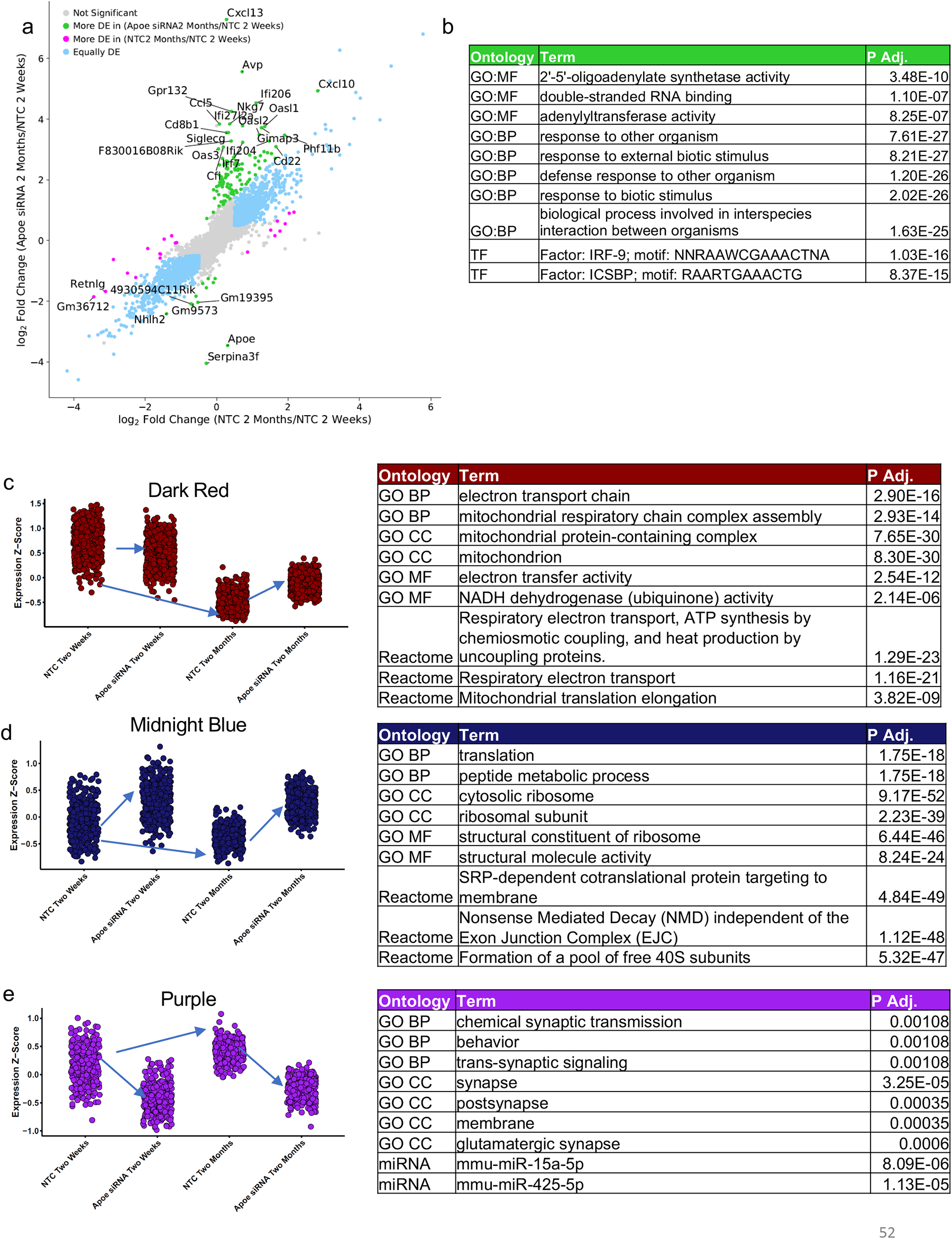
*ApoE* silencing causes sustained activation of the immune response. (**a**) Comparison of differential expression after silencing *ApoE* or controls (NTC) for two months versus two-week control (NTC) treatment shows the effect of *ApoE* silencing on the transcriptome over the course of disease progression in 5XFAD mice. (**b**) Gene ontology showing upregulated pathways over time (2 weeks and 2-months) after *ApoE* silencing. (**e-g**) WGCNA module gene average expression z-score and gene ontology results for modules that are significantly correlated with amyloid burden and treatment conditions (R > 0.3, p-value < 0.05). 5xFAD mouse model; Age: 11 and 17 weeks old. Timepoints: 2-weeks post treatment, 2-months post injection. N=9-10 per group at 2-weeks post injection; n=18-20 per group 2-months post injection.

### Gene network analysis identifies clusters associated with *ApoE* silencing over time

Over time, *ApoE* silencing normalized the expression of 21 genes whose expression changed during disease progression and altered the expression of 129 genes (**Fig. 5a, b**). Figure 5b shows upregulated GO pathways. To capture how *ApoE* silencing alters disease-associated gene sets, we performed weighted gene co-expression network analysis (WGCNA) (Androvic et al., 2020). We identified WGCNA modules that positively or negatively correlated with disease progression (p-value < 0.05, absolute value of R > 0.3) (**Extended Data Fig. 8e**) and looked more closely at modules that were also significantly changed after silencing of *ApoE*. The dark red and midnight blue modules were negatively correlated with amyloid burden and rescued by *ApoE* silencing (**Fig. 5c-d**). The purple module was positively correlated with amyloid burden and downregulated by *ApoE* silencing (**Fig. 5e**). These three modules were enriched for mitochondrial and electron transport chain genes (dark red), ribosomal subunit proteins, translation, and translational regulation (midnight blue), and synapse-related genes, including glutamatergic synapses, and predicted targets of mmu-miR-15a-5p and mmu-miR-425-5p (purple module) (**Extended Data File 6-7**). The gene ontologies associated with ApoE silencing are like those of ApoE2 carriers, who have increased protection against AD (Hernandez-Ortega et al., 2016).

## Discussion

The presence of *ApoE4* is the most significant risk factor for developing late onset AD (E. H. Corder, 1993; Hardy and Escott-Price, 2019; Yu et al., 2014). Some *ApoE*-based therapeutics, including an *ApoE* mimetic and AAV gene delivery of *ApoE2,* are in Phase 1 clinical trials (Williams et al., 2020), but many previously evaluated *ApoE* therapies (e.g., Bexarotene and Rosiglitazone) have been ineffective in humans, and none silence *ApoE4* to the levels seen in transgenic proof-of-concept studies (Williams *et al*., 2020; Yamazaki et al., 2019). Here, we achieve potent, CNS-selective silencing of *ApoE* using therapeutic siRNAs. Silencing CNS *ApoE*, even after the onset of pathologic changes, improves neuropathology in adult mice without disrupting systemic cholesterol homeostasis. <u>Our results show a multi-faceted role for ApoE in the progression of AD and propagation of amyloid pathology. Indeed, when evaluating the literature, ApoE has been implicated in what seems like endless mechanisms and cellular processes. We show that silencing *ApoE* with therapeutic siRNAs improves amyloid pathology via reduction in amyloid fragment production and reduction of ApoE rich cores present for plaque formation, which further allows for effective neuroprotective microglial clearance.</u> These results challenge prior conclusions that therapeutic silencing of *ApoE* is not effective in the adult brain and support a gain of function impact of ApoE expression on toxic neuroinflammation.

Building on previous studies, we found no detectable trafficking of *ApoE* in and out of the adult CNS, even after the removal of systemic ApoE (Huynh *et al*., 2019; Lane-Donovan *et al*., 2016). Silencing liver *ApoE* did not alter Aβ pathology in APP/PSEN1 mice, suggesting that systemic *ApoE* does not exacerbate disease pathology, consistent with previous reports (Huynh *et al*., 2019). Future studies could evaluate the effects of long-term silencing of liver *ApoE* on AD pathogenesis.

Importantly, ours is the first study to show that silencing *ApoE* with siRNAs can be therapeutic: silencing *ApoE* after the onset of cellular changes significantly reduced amyloid pathology in 5xFAD mice. Although a reduction in Aβ plaques correlates well with behavioral improvements (Janus et al., 2015), we did not investigate the impact of silencing *ApoE* on AD-related cognitive dysfunction in these experiments, because the 5xFAD nor APP/PSEN1 models do not develop cognitive deficits until at least 6 months of age, and the results can be highly variable (Devi and Ohno, 2012; Savonenko et al., 2005). We did, however, confirm that silencing *ApoE* with di-siRNA for extended periods does not negatively affect cognition in young-adult 5xFAD mice, and thus, appears to be well-tolerated.

Mechanistically, our findings suggest that ApoE modulates APP processing, serves as a scaffold for amyloid oligomerization that blocks pattern recognition receptors, and prevents immune-mediated amyloid degradation, which leads to microglial dysfunction and cellular death (Fitz *et al*., 2020; Sala Frigerio *et al*., 2019; Ulrich *et al*., 2018). The ApoE-dependent reduction in amyloid CTFs was surprising, particularly as there was no effect on expression of APP or APP processing machinery (APP, BACE1, PSEN). Either CTFs are degraded alongside amyloid fragments or silencing *ApoE* reduces CTF production. Changes in production could be due to *ApoE*-dependent changes in APP membrane localization and cellular trafficking (Wang et al., 2021; Ye et al., 2005). Future studies are needed to determine how *ApoE* silencing reduces CTF burden in vivo.

We show that silencing CNS *ApoE* reduces *ApoE*-rich plaque cores, causing structural changes in amyloid plaques. We hypothesize that reduction of *ApoE* cores slows the formation of amyloid plaques and allows for effective immune-related clearance of toxic amyloid and pathologic improvement. Disease pathology drives *ApoE* expression in activated microglia, which may both promote plaque formation via post-translational modification and enhance disease associated inflammation (Sala Frigerio *et al*., 2019). We also found that *ApoE* also serves as a key modulator of immune response pathways in AD, promoting the expression of AD-related genes and inflammation, consistent with previous reports (Krasemann et al., 2017; Sala Frigerio *et al*., 2019; Ulrich *et al*., 2018; Wan et al., 2020). Although we did not perform cell-type-specific RNAseq, expression signatures after silencing *ApoE* resembled those seen in DAM cells. Microglial responses in AD are dynamic, with DAM signatures being associated with acute, clearance-related inflammation, as well as chronic, harmful inflammation (Sarlus and Heneka, 2017). DAM have been implicated in acute inflammation required to acutely clear toxic pathogens, as well as chronic, possibly harmful, inflammatory processes (Sarlus and Heneka, 2017). Given their dynamic nature, it is difficult to draw conclusions regarding the implications of DAM expression signatures, and at what point the switch between acute and chronic activation occurs. Whether this corresponds with protective acute immune responses or chronic inflammatory signaling warrants further investigation (Pascoal et al., 2021; Sala Frigerio *et al*., 2019). We hypothesize that silencing *ApoE* slows the turning point from acute to chronic neuroinflammation in the context of AD (Friker et al., 2020).

The role of *ApoE* in AD has been muddled by differences in conferred risk between populations (i.e., Asian, European, African American) (Farrer and Neil Risch, 1997; M. X. Tang, 1996; Osuntokun BO, 1995; Reitz and Mayeux, 2014), and confounded by factors like BMI and diabetes. GWAS studies point towards a protective role of ApoE2 (Reiman et al., 2020) and *in vivo* analyses suggest a gain-of-function toxicity for ApoE4 (Arboleda-Velasquez et al., 2019; Reiman *et al*., 2020). In support of the gain-of-function model, individuals heterozygous for ApoE4 (e.g., ApoE2/4 or ApoE3/4) have similar rates of Aβ burden, suggesting that the toxicity of ApoE4 outweighs a protective role for ApoE2 (Yamazaki *et al*., 2019). Although evidence also suggests a role for ApoE4 loss-of-function in AD, this model is less convincing in the context of late-onset AD. For example, increasing the expression of ApoE4 in AD mouse models does not reduce Aβ deposition (Hu et al., 2015; Hudry et al., 2013), and a dose-dependent relationship between ApoE4 levels and amyloid pathology exists in humans (Reiman et al., 2009). Our results confirm a gain-of-function effect of ApoE on both amyloid plaque formation and toxic immune responses. Moreover, our comparison with published datasets demonstrates similar expression profiles between ApoE2 presence and silencing *ApoE*, supporting *ApoE* silencing as a therapeutic approach for AD.

We effectively evaluate the spatial and functional relationships between systemic and CNS pools of *ApoE*, the impact of these pools on AD pathological phenotypes, and identify potential mechanisms through which *ApoE* impacts AD pathology. With a single administration of 237 µg di-siRNA, we achieved potent, sustained silencing of CNS *ApoE*, likely due to the integration of a fully-modified scaffold that allows the siRNA to be internalized by the cell and stored in the endosome for sustained release (Brown et al., 2020). Moving forward, this study provides the rationale and technology necessary to further evaluate the relative impact of therapeutic *ApoE* silencing on other neurodegenerative diseases, including Huntington’s disease and tauopathies (Litvinchuk *et al*., 2021), and to further explore *ApoE* modulation as a therapeutic paradigm for AD. The therapeutic siRNAs developed in this study can be taken directly to clinical trials, accelerating development of much needed disease-modifying therapies.

## Online Methods

### Oligonucleotide synthesis

Oligonucleotides were synthesized using modified (2ʹ-F, 2ʹ-O-Me, LNA) phosphoramidites with standard protecting groups. Phosphoarmidite solid-phase synthesis was done on a MerMade12 (BioAutomation) using modified protocols. Unconjugated oligonucleotides were synthesized on controlled pore glass (CPG) functionalized with a long-chain alkyl amine (LCAA) and unylinker terminus (Chemgenes). GalNAc-conjugated oligonucleotides were grown on custom 3’GalNAc-CPG (Sharma et al., 2018), Divalent oligonucleotides (DIO) were synthesized on modified solid support (Alterman *et al*., 2019), and Vinyl-phosphonate (VP) phosphoramidite was synthesized as described. Phosphoramidites were prepared at 0.1 M in anhydrous acetonitrile (ACN), with added dry 15% dimethylformamide (DMF) in the 2’-OMe U amidite. 5-(Benzylthio)-1H-tetrazole (BTT) was used as the activator at 0.25 M. Detritylations were performed using 3% trichloroacetic acid in dichloromethane (DCM). Capping was done with non-tetrahydrofuran-containing reagents CAP A, 20% n-methylimidazole in ACN and CAP B, 20% acetic anhydride (Ac2O), and 30% 2,6-lutidine in ACN (Synthesis reagents were purchased at AIC). Sulfurization was performed with 0.1 M solution of 3-[(dimethylaminomethylene)amino]-3H-1,2,4-dithiazole-5-thione (DDTT) in pyridine (ChemGenes) for 3 min. Phosphoramidite coupling times were 3 min for all amidites used.

### Vinyl phosphonate deprotection

The VP-containing oligonucleotides, still on solid support, were treated post synthesis with an anhydrous mixture of trimethylsilyl bromide/ACN/DMF/pyridine (3:9:9:1) for 1 h at room temperature with gentle agitation. The reaction was then quenched with water and the CPG was then rinsed with, ACN, DCM and allowed to dry, before being deprotected normally as described below.

### Deprotection and purification of oligonucleotides

Divalent and conjugated oligonucleotides (DIO, GalNAc) were cleaved and deprotected with standard conditions using aqueous ammonia at 55⁰C for 16h. VP-containing oligonucleotides were cleaved and deprotected as described (O’Shea et al., 2018). Briefly, CPG-containing VP-oligonucleotides were treated with a solution of 3% Diethylamine (DEA) in aqueous ammonia at 35⁰C for 20h. The solutions containing deprotected oligonucleotides were filtered to remove the CPG, and dried under vacuum in Speed-vac. The resulting pellets were re-suspended in 5% ACN in water. Purification was performed on an Agilent 1290 Infinity II HPLC system, equipped with a Source 15Q anion exchange column (GE Healthcare) using the following conditions: eluent A, 20% ACN, 20 mM sodium acetate pH 7; eluent B, 1 M sodium perchlorate in 20% ACN; gradient, 10% B 3 min to 35% B 18 min, at 60 °C. Peaks were monitored at 260 nm. Pure fractions were collected and dried in Speed-vac. Oligonucleotides were re-suspended in 5% ACN and desalted through fine Sephadex G-25 media (GE Healthcare), and lyophilized.

### LC–MS analysis of oligonucleotides

The identity of oligonucleotides were verified by LC–MS analysis on an Agilent 6530 accurate mass Q-TOF using the following conditions: buffer A: 100 mM hexafluoroisopropanol (HFIP)/ 9 mM triethylamine (TEA) in LC–MS grade water; buffer B:100 mM HFIP/9 mM TEA in LC–MS grade methanol; column, Agilent AdvanceBio oligonucleotides C18; gradient 0% B 1 min, 0–40% B 8 min, temperature, 45°C; flow rate, 0.5 ml/min. LC peaks were monitored on UV (260 nm). MS parameters: source, electrospray ionization; ion polarity, negative mode; range, 100–3,200 m/z; scan rate, 2 spectra/s; capillary voltage, 4,000; fragmentor, 180 V. Reagents were purchased from Fisher Scientific, Sigma Aldrich and Oakwood Chemicals, and used per manufacturer’s instructions, unless otherwise stated.

### Animal Studies

All experimental studies involving animals were approved by the University of Massachusetts Medical School Institutional Animal Care and Use Committee (IACUC Protocols #A-2411 and #A-1744) and performed according to the guidelines and regulations therein described. APP/PSEN1 (MMRRC 34832-JAX) mice were bred in house and obtained from Jackson Lab. 5xFAD mice (MMRRC Stock No: 34840-JAX) were obtained from Jackson Lab (Experiments were performed on hemizygote APP/PSEN1 and 5xFAD mice). Initial aged APP/PSEN1 mouse colonies were a gift from Dr. Arya Biragyn from NIA.

### Stereotactic ICV injections

10 μl di-siRNA was administered bilaterally (5 uls per ventricle) into the lateral ventricles of mice as previously described (Alterman et al., 2019). Briefly, mice were anesthetized using avertin and prepared using standard aseptic technique. Stereotaxic devices were using to hold injection needles and identify injection location. After the identification of the bregma, the needle was placed 1 mm laterally, 0.2 mm posterior, and 2.5 mm caudally. Injection was performed at 500 nl/min. Mice were then monitored until fully sternal.

### Sub-cutaneous injections

GalNAc-conjugated siRNAs were injected subcutaneously (SC) into mice. Each animal received a 10 mg/kg dose in 200 ul volume. For anti-siRNAs, mice received 1 mg/kg SC.

### Tissue collection

mRNA quantification was performed as described in Alterman et al., 2015. Briefly, tissue punches were stored in RNAlater (Invitrogen #AM7020) and homogenized in Quantigene 2.0 homogenizing buffer (Invitrogen, QG0517) with proteinase K (Invitrogen, 25530-049). mRNA was detected according to the Quantigene 2.0 protocol using the following probe sets: mouse HPRT (SB-15463), mouse PPIB (SB-10002), mouse apoE (SB-13611).

### Protein quantification

For analysis of ApoE protein expression in mouse brain samples, WES by ProteinSimple was used as previously described (Alterman *et al*., 2019) and (Gentalen E.T., 2015). Briefly, tissue punches were collected as above and flash-frozen and placed at -80°C. After addition of RIPA buffer with protease inhibitors, samples were homogenized and stored at -80°C. Protein amount was determined using Bradford Assay. Samples were diluted in 0.1x sample buffer (ProteinSimple) to ∼0.2-0.4 μg/μl. Anti-ApoE antibody (Abcam, 183597) was diluted 1:200 in antibody diluent (ProteinSimple) and loading control, anti-Vinculin (Invitrogen, 700062), was diluted 1:1000 in antibody diluent. Assay was performed as described by ProteinSimple protocol using the 16-230 kDa plate (SM-W004). The separation electrophoresis and immunodetection are performed automatically in the capillary system using the default system settings. Once loaded, the separation electrophoresis and was performed automatically. Results were analyzed using the Compass for Protein Simple software. APP, CTFs, and AB42 protein was evaluated using traditional western blots as described below: In brief, snap-frozen hippocampal and cortical brain samples were extracted in PBS, 1 mM EDTA, 1 mM EGTA, 3 µl/ml protease inhibitor mix (Sigma, Munich, Germany) as previously described (Venegas et al., 2017). Homogenates were extracted in RIPA buffer (25 mM Tris-HCl, pH 7.5, 150 mM NaCl, 1% NP40, 0.5% NaDOC, 0.1% SDS), centrifuged at 100,000 x g for 30 min and the pellet containing insoluble Aβ was solubilized in 2% SDS, 25 mM Tris-HCl, pH 7.5. In addition, the SDS-insoluble pellet was extracted with 70% formic acid in water. Formic acid was removed using a speed vac (Eppendorf, Hamburg, Germany) and the resulting pellet was solubilized in 200 mM Tris-HCl, pH 7.5. Samples were separated by 4-12% NuPAGE (Invitrogen, Karlsruhe, Germany) using MES or MOPS buffer and transferred to nitrocellulose membranes. APP and Aβ were detected using antibody 6E10 (Covance, Münster, Germany) and the c-terminal APP antibody 140 (CT15). α-tubulin served as housekeeping control. Immunoreactivity was detected by enhanced chemiluminescence reaction (Millipore, Darmstadt, Germany) or near-infrared detection (Odyssey, LI-COR). Chemiluminescence intensities were analyzed using Chemidoc XRS documentation system (Biorad, Munich, Germany).

### Cholesterol

Serum cholesterol was measured using the Abcam LDL and HDL cholesterol quantification kit (ab65390). LDL and HDL were separated using the included precipitation buffer that uses a water-soluble non-ionic polymer to precipitate the fractions (Liu et al., 2012). The assay uses cholesterol esterase to hydrolyze cholesteryl ester into free cholesterol. Next, cholesterol oxidase acts on free cholesterol and to produce a color at 570 nm that is proportional to the amount of cholesterol in the sample. Briefly, serum was collected prior to euthanasia. 2x buffer was added to 50 μl of serum and incubated at room temperature for 10 minutes. Samples were spun for 10 minutes at 2000 rpm and the supernatant (LDL fraction) was placed in a separate tube. The pellet (HDL fraction) was resuspended in 200 ul PBS. The samples were diluted, and cholesterol levels analyzed according to the package instructions.

### Histology and analysis

For evaluation of amyloid plaque burden, four 40-uM formalin-fixed brain slices per animal were stained in a free-floating format with either X-34 dye (Sigma-Aldrich, #SML1954), APP6E10 (BioLegend, #803001), or AB42 (Cell Signaling, D9A3A). For X-34 staining, formalin-fixed brain slices were washed in PBS for 5 minutes, incubated in 100uM X-34 (dissolved in 40% EtOH) for 10 minutes, rinsed 5x in water, incubated in 0.2% NaOH in 80% EtOH for 2 minutes, then washed in distilled water for 10 minutes, and mounted for imaging. Images were performed with a 20x objective on Leica inverted microscope. Number and size (area, volume) was quantified using ImageJ thresholding and 3D object counter and normalized to controls. For IHC, three 40-μM slices were placed in 70% formic acid for 15 minutes, washed three times in PBS, placed in 1% H2O2 for 30 minutes, washed three times in PBS, blocked and permeabilized in 10% blocking serum and 0.5% Triton-X. Samples were placed in biotin-labeled APP 6E10 (BioLegend #803007) at 4°C overnight and developed using ABC (VectorLab #PK-6200) DAB peroxidase system (VectorLab #SK-4100). Samples were allowed to dry and dehydrated in ethanol, cleared with xylenes and cover-slipped. Images were acquired with a 5x objective using tiling. Analysis was performed using ImageJ. Briefly, images were converted to 8-bit images. Thresholding was applied to measure number and size of amyloid plaques. Amyloid burden was reported as percent of total area measured. Values across at least three slices per animal were averaged and compared to controls.

For immunofluorescence, slices were treated with 70% formic acid for 15 minutes, incubated in blocking buffer (PBS with 1% BSA and 0.3% Triton X-100) for 1 hour at room temperature, incubated with primary antibodies overnight at rotating 4 °C (APP6E10: 1:1000, AB-42: 1:200, IBA1: 1:500, ApoE: 1:500, GFAP 1:500), washed with PBS, incubated with Alexa-Fluor-labeled secondary antibodies diluted 1:500 in blocking buffer for 2 hours at room temperature, washed with PBS, and mounted with ProLong Diamond. Imaging was performed on Leica Inverted microscope.

### Microglial clustering

Samples were processed for immunofluorescence as described above with IBA1 primary antibody (ab178846, 1:500) and Alexa-Fluor-labeled secondary antibodies diluted 1:500. Five random regions of interest per animal were acquired from the cortex using a Leica Inverted microscope. The number of free and clustered (plaque associated) microglia per field of view was quantified using ImageJ. The numbers were averaged per animal.

### Aβ ELISAs

For APP/PSEN1 mice, Aβ-42 (ThermoFisher, #KHB3544) and Aβ-40 (ThermoFisher, #KHB3481) were measured using ELISAs. For 5xFAD mice, Aβ-42, Aβ-40, and Aβ-38 were quantified using multiplex MSD plates (MSD, #K15200E). Brain homogenates were solubilized in RIPA buffer with protease inhibitors and spun down for 20 minutes. The RIPA fraction was transferred to a new tube. The pellet was resuspended in 70% formic acid and incubated rotating for 2 hours. Samples were then spun down and the supernatant transferred to a new tube and used for ELISA assays. ELISAs and MSD assays were performed and analyzed according to the manufacturer’s instructions.

### RNA sequencing and data analysis

*Library Preparation:* RNA from brain punches was purified using the NEB RNA purification kit (NEB, #T2010S). RNA libraries were prepared using TruSeq kit (Illumina, #20020595). Samples were run on HiSeq Flow 4000 for NextSeq 550 single end with read lengths of 100 and 72, respectively. *Mapping:* Reads were then mapped to a custom mm10 genome containing human APP for the APP/PSEN1 mice, a custom genome containing human APP and PSEN1 for the 5XFAD mice, and a custom genome of mm10 containing human APP and APOE4 for the 3XAD mice, using STAR v2.7.6a (Dobin et al., 2013) and transcript expression was calculated with RSEM v1.3.3 (Dewey, 2011). Genes with an estimated count of less than 5 in at least half the samples were removed from further analysis. *Differential Expression:* Differential expression was calculated with DESeq2 v1.22.2 using an alpha of 0.05 and apeglm log2 fold change shrinkage (Love et al., 2014; Zhu et al., 2019). The presence of siRNA seed complementarity (7-mer complementary to siRNA guide strand positions 2-8) in all genes with annotated 3′-UTRs in Ensembl GRMCm38.p6 annotations was determined with a custom Python script. siRNA seed enrichment in downregulated vs unchanged and upregulated transcripts was calculated using a Fisher Exact Test. Data was plotted using Matplotlib. *Gene Set Enrichment Analysis:* Gene set enrichment analysis was conducted using the fgsea R package (Korotkevich et al., 2016) and plotted using the viseago R package (Brionne et al., 2019), with custom settings for heatmap color to indicate if a gene set is upregulated or downregulated. *WGCNA:* Data for WGCNA (Langfelder and Horvath, 2008) were first filtered to remove genes with 10 or fewer reads in any sample, batch effect corrected for sex using the ComBat-Seq R package (Zhang et al., 2020) and transformed using the vst function in DESeq2. WGCNA with additional k-means clustering was conducted using the CoExpNets R package (Botía et al., 2017) with default settings. Gene ontology of modules was performed with gprofiler2 (Kolberg et al., 2020). *Disease Associated Microglia and Astrocyte Subtype Analysis:* Marker genes for disease associated microglia (DAM) and inflammatory astrocyte subtypes were taken from previously published single cell RNA sequencing data (Fitz *et al*., 2021; Grubman et al., 2021; Hasel *et al*., 2021; Mathys *et al*., 2017). A custom GMT file was created using these marker genes and GSEA was performed with this GMT file with GSEAPY, using 1000 permutations. *Transcription Factor Enrichment Analysis:* Transcription factor enrichment analysis of upregulated genes was performed using the MAGIC 1.1 python package, using the “1kb_gene” matrix (Roopra, 2020). *Single-Cell RNA Sequencing Analysis:* Analysis of single-cell RNA sequencing data from (Sala Frigerio *et al*., 2019) was performed according to their methods. Briefly, .loom files for microglia from APP/PSEN1 mice and APP-NL-G-F mice were downloaded from scope.bdslab.org, gene expression matrices were extracted, and cells were clustered using Seurat 4.0.5 (Butler et al., 2018). For the APP-NL-G-F microglia, PCA was performed on the 4,967 most variable genes and the first 10 PCs were used for cluster analysis. For the APP/PSEN1 microglia, PCA was performed on the 4,777 most variable genes and the first 14 PCs were used for cluster analysis. In both cases, a resolution of 0.25 was used for the *FindClusters* command and the data was visualized using Seurat’s t-SNE and *FeaturePlot* implementations. Marker genes for each cluster were found using the *FindAllMarkers* command with only.pos=TRUE. Genes with an avg_log2FC of >0.5 and p_val_adj<0.05 were used as marker genes. The cluster marker genes were then cross-referenced with the clusters from (Sala Frigerio *et al*., 2019).

### Statistics

Each n represents an independent biological sample. All graphs show means ± S.D. All statistics were performed using Prism GraphPad v.8, using either two-tailed, unpaired t-tests or one-way ANOVA with Dunnett correction. Box plots: center line, median; top of box, 75% quartile; bottom of box, 25% quartile; whiskers, minimum and maximum. The GO enrichment analysis was performed according to Fisher’s exact test followed by a Bonferroni multiple comparison correction.

## Reporting summary

Further information on research design is available in the reporting summary linked to this article.

## Data availability

All data generated during this study are either included in this article or are available from the corresponding author on reasonable request. Raw and processed RNA sequencing data was uploaded to the Gene Expression Omnibus (GSE186756).

Custom scripts used for RNA sequencing analysis have been deposited to https://github.com/shildebr12/Ferguson_et_al_2021.

## Author contributions

C.M.F, A.K., and E.R. conceived of the project. C.M.F. and A.K. contributed to experimental design. C.M.F, B.M.D.C.G., and A.C., performed ICV injections. C.M.F. and A.G. performed mouse breeding. C.M.F, J.B, F.S., and M.T.H performed sample preparation and analysis. S.H. performed RNA seq analysis. D.E., M.R.H., L.V., J.S., and N.M. synthesized compounds. C.M.F. and A.K. wrote the manuscript.

## Acknowledgements

This work was supported by NS 104022, NID GM-13 183901, and ADDF 20170101/Rogaev; National Research Service Award 1F30AG066373-01, and T32GM107000. We would like to thank Dr. Arya Biragyn, NIA, for gifting us AD mouse strains, and the Animal Medicine Group at UMass Chan Medical School. We would like to thank Emily Haberlin and Darryl Conte for editing the manuscript.

---

**Extended Data Figure 1: Reduction of liver ApoE has no detectable impact on AD neuropathology in APP/PSEN1 animal model but causes significant increase in serum LDL-cholesterol.** (**a, b**) Experimental key and timeline. *ApoE* mRNA (**c**) and (**d**) protein expression in the hippocampus, cortex, and liver 2-months post administration of control or GalNAc-siRNA^APOE^. (**e**) Serum HDL and LDL cholesterol in controls and treated mice. (**f**) Amyloid burden (APP6E10) in the cortex 2-months post injection in control (GalNAc-siRNA^NTC^) and treated animals (GalNAc-siRNA^APOE^). (**g**) Sex-stratified quantification of APP6E10-positive cortex plaques. (**h**) Insoluble Aβ-42 in cortex samples. (**i**) Formic acid soluble Aβ-42 in cortex samples. GalNAc-siRNA^NTC^: n=3 males; GalNAc-siRNA^APOE^: n=6 males; Dose = 10 mg/kg). Statistical analysis: one-way ANOVA or T-test. Error bars are SD. (****p <0.0001; *** p<0.002; ** p<0.01; * p<0.05).

**Extended Figure 2: Liver and CNS *ApoE* protein do not traffic between CNS and Liver in APP/PSEN1 mice. (a)** Raw western blots showing ApoE protein in the hippocampus, cortex, and liver (top to bottom) after administration of di-siRNA^NTC^ or di-siRNA^APOE^. (**b**) Raw western blots showing ApoE protein in the liver but not the hippocampus or cortex after administration of GalNAc^NTC^ or GalNAc^APOE^. ApoE 37 kDa, bottom normalized to loading Vinculin, 116 kDa; top). Protein evaluated with WES for Protein Simple.

**Extended Data Figure 3: Silencing ApoE with di-siRNA^APOE^ for 2-months does not affect cognitive function in 5xFAD mice. (a)** Study design. Day 1: open field testing. Day 2: Novel Object Recognition. Day 3: y-maze. (**b**) Time spent in periphery (left), distance traveled (middle), and average velocity (right) during open field testing. (**c**) Discrimination ratio (left), distance traveled (middle), and average velocity (right) during novel object recognition. (**d)** Spontaneous alternation (left), distance traveled (middle), and average velocity (right) during open field testing. Experiments performed and analyzed using Noldus EthosVision XT15 software. Statistics performed using GraphPad. T-tests per assay, sex separated. *: p<0.0332; ** p<0.0021; *** p<0.0002; **** p<0.0001. n=9-10 per sex per group. Timepoint: 16-weeks-old, 2 months post injection. Di-siRNA^NTC^ shown in grey, and di-siRNA^ApoE^ shown in green.

**Extended Data Figure 4: Short-term Silencing of *ApoE* does not reduce Aβ burden.** (**a**) Immunohistochemistry showing amyloid burden (APP 6E10) in control (di-siRNA^NTC^; top) and ApoE silenced (di-siRNA^APOE^; bottom) 5xFAD animals. Scale bar: 1000 μm (**b**) High-resolution images showing onset of plaque deposition in the hippocampus and cortex. Scale bar: 500 μm. (**c**) Quantification of amyloid burden (top), amyloid plaque size (middle), and amyloid plaque number (bottom). Age: 11 weeks; 2 weeks post injection. Female: NTC: n=4/group, APOE: n=5/group; Male: NTC: n=5/group, APOE: n=5/group. Statistics: GraphPad Prism: t-tests per brain region. *: p<0.0332; ** p<0.0021; *** p<0.0002; **** p<0.0001. Di-siRNA^NTC^ shown in grey, and di-siRNA^APOE^ shown in green.

**Extended Data Figure 5: Silencing *ApoE* primarily reduces plaque number not size. (a)** Amyloid plaque size and (**b**) Amyloid plaque number measured using APP6E10 antibody. (**c**) Aβ-42 plaque size and (**d**) Aβ-42 plaque number measured using Aβ-42-specific antibody (D9A3A) in 5xFAD treated and control mice, separated by sex. (**e**) MSD ELIDA quantification of Aβ-42 and Aβ-40 in soluble brain fractions. Statistics: t-test per brain region; sex separated. *: p<0.0332; ** p<0.0021; *** p<0.0002; **** p<0.0001. n=9-10 per sex per group. Timepoint: 17 weeks old, 2 months post injection. Di-siRNA^NTC^ shown in grey, and di-siRNA^ApoE^ shown in green.

**Extended Data Figure 6: Raw western blot files for analysis of APP, CTFs, and Aβ42 in 5xFAD mice.** (**a**), (**b**) Raw western blots of cortex in a subset of 5xFAD samples. (**c**), (**d**) Raw western blots of hippocampus in a subset of 5xFAD samples. The last lane on all blots is a repeated internal control sample. N=4/5 per group.

**Extended Data Figure 7: Confirmation of RNA expression using RTqPCR.** Statistics: One-way ANOVA. *: p<0.0332; ** p<0.0021; *** p<0.0002; **** p<0.0001. n=10-20 per group. Di-siRNA^NTC^ shown in grey, and di-siRNA^ApoE^ shown in green.

**Extended Data Figure 8: Further analysis of RNA Sequencing: (a-c**) Mean expression z-scores for genes that are upregulated with *ApoE* knockdown at both timepoints (**a**), 2-weeks (**b**) or 2 months (**c**) post-treatment. (**d**) Comparison of differentially expressed genes in male and female 5XFAD mice at 2 months post-treatment. (**e**) Correlation between WGCNA clusters and either treatment or amyloid burden.

**Extended Data Figure 9: (a)** Comparison of gene overlap between *ApoE* silencing for 2-months in 5XFAD and marker genes of disease associated microglia (Mathys et al., 2017) (**b**), activated and interferon response microglia subtypes (Sala Frigerio et. al. 2019), and (**d**) plaque associated microglia, (**e**) microglial responses after Aβ/APOE3 or Aβ/APOE4 injection (Fitz et al., 2021), and (**f**) marker genes of astrocytes (Hasel et al., 2021).

**Extended Data Table 1: Detailed sequence and chemical modification patterns of siRNAs.** Chemical modifications are designated as follows, “#” –phosphorothioate bond, “m” – 2’-O-Methyl, “f” – 2’-Fluoro, “P” – 5’ Phosphate, “V” – 5’-(*E*)-Vinylphosphonate. “L” – LNA. “DIO” – di-siRNA.

**Extended Data Taable 2:** Markers of Liver toxicity in APP/PSEN1 mice

**Extended Data Table 3: Statistical Differences in Amyloid Plaque Burden, Number, and Size (APP6E10) 5xFAD mice, 2-months post injection (% area).** Statistics Performed using Graph Pad Prism: T-test per brain region (sex separated). N=9-10 per sex per group.

**Extended Data Table 4: Statistical Differences in Amyloid Plaque Burden, Number, and size (Aβ-42) 5xFAD mice, 2-months post injection (% area).** Statistics Performed using Graph Pad Prism: T-test per brain region (sex separated). N=9-10 per sex per group.

**Extended Data File 5: RNAseq Differential Expression Tables Extended Data File 6: GSEA Results**

**Extended Data File 7: WGCNA Results**

**Extended Data File 8: APOE2 Vs APOE4 Human GSEA**

