## Supplemental Figures for "Silencing of *ApoE* with Divalent siRNAs Drives Activation of Immune Clearance Pathways and Improves Amyloid Pathology in Mouse Models of Alzheimer’s Disease"

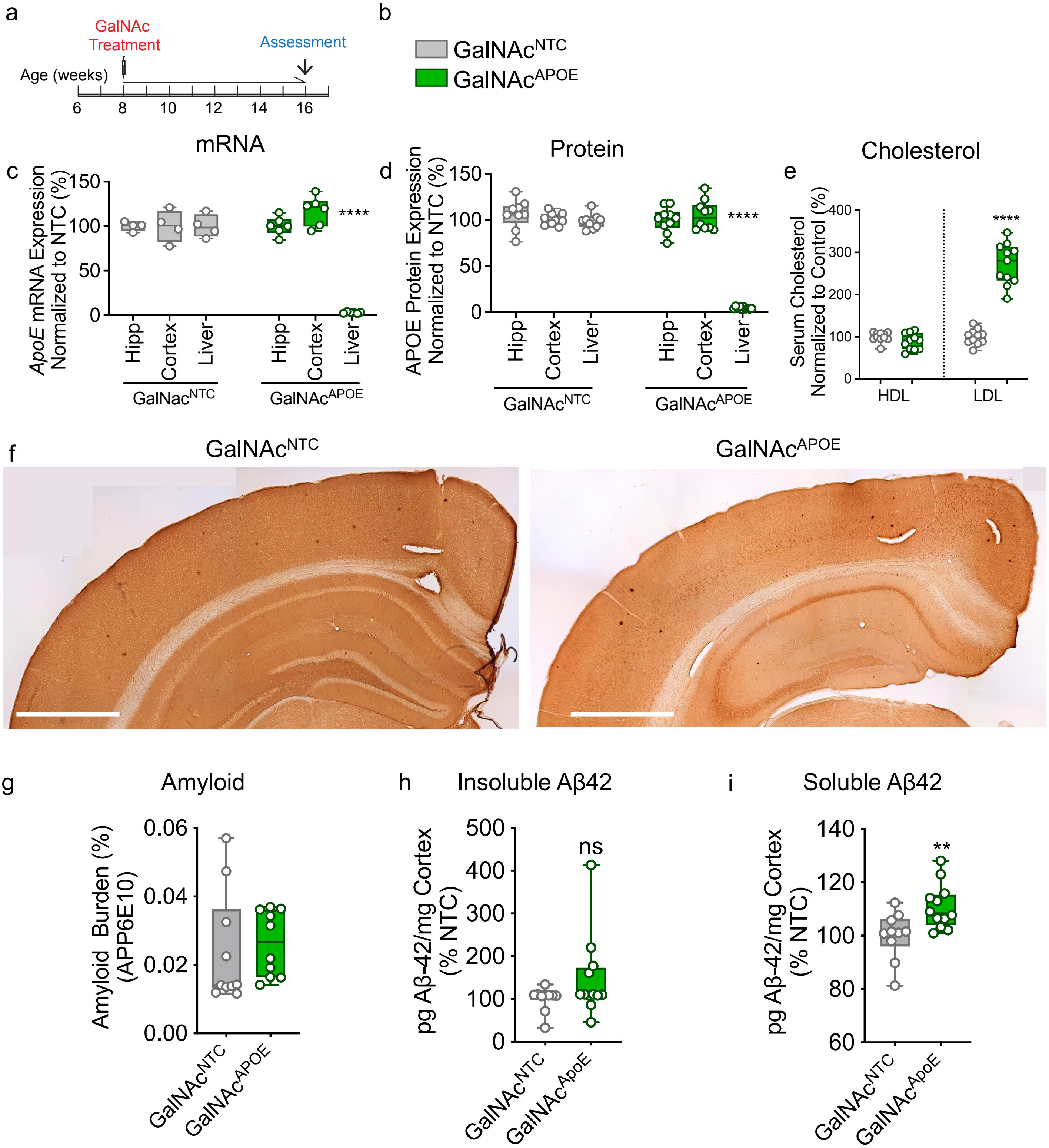


**Extended Data Figure 1: Reduction of liver ApoE has no detectable impact on AD neuropathology in APP/PSEN1 animal model but causes significant increase in serum LDL-cholesterol.** (**a**, **b**) Experimental key and timeline. *ApoE* mRNA (**c**) and (**d**) protein expression in the hippocampus, cortex, and liver 2-months post administration of control or GalNAc-siRNA$^{APOE}$. (**e**) Serum HDL and LDL cholesterol in controls and treated mice. (**f**) Amyloid burden (APP6E10) in the cortex 2-months post injection in control (GalNAc-siRNA$^{NTC}$) and treated animals (GalNAc-siRNA$^{APOE}$). (**g**) Sex-stratified quantification of APP6E10-positive cortex plaques. (**h**) Insoluble Aβ-42 in cortex samples. (**i**) Formic acid soluble Aβ-42 in cortex samples. GalNAc-siRNA$^{NTC}$: n=3 males; GalNAc-siRNA$^{APOE}$: n=6 males; Dose = 10 mg/kg). Statistical analysis: one-way ANOVA or T-test. Error bars are SD. (****p <0.0001; *** p<0.002; ** p<0.01; * p<0.05).

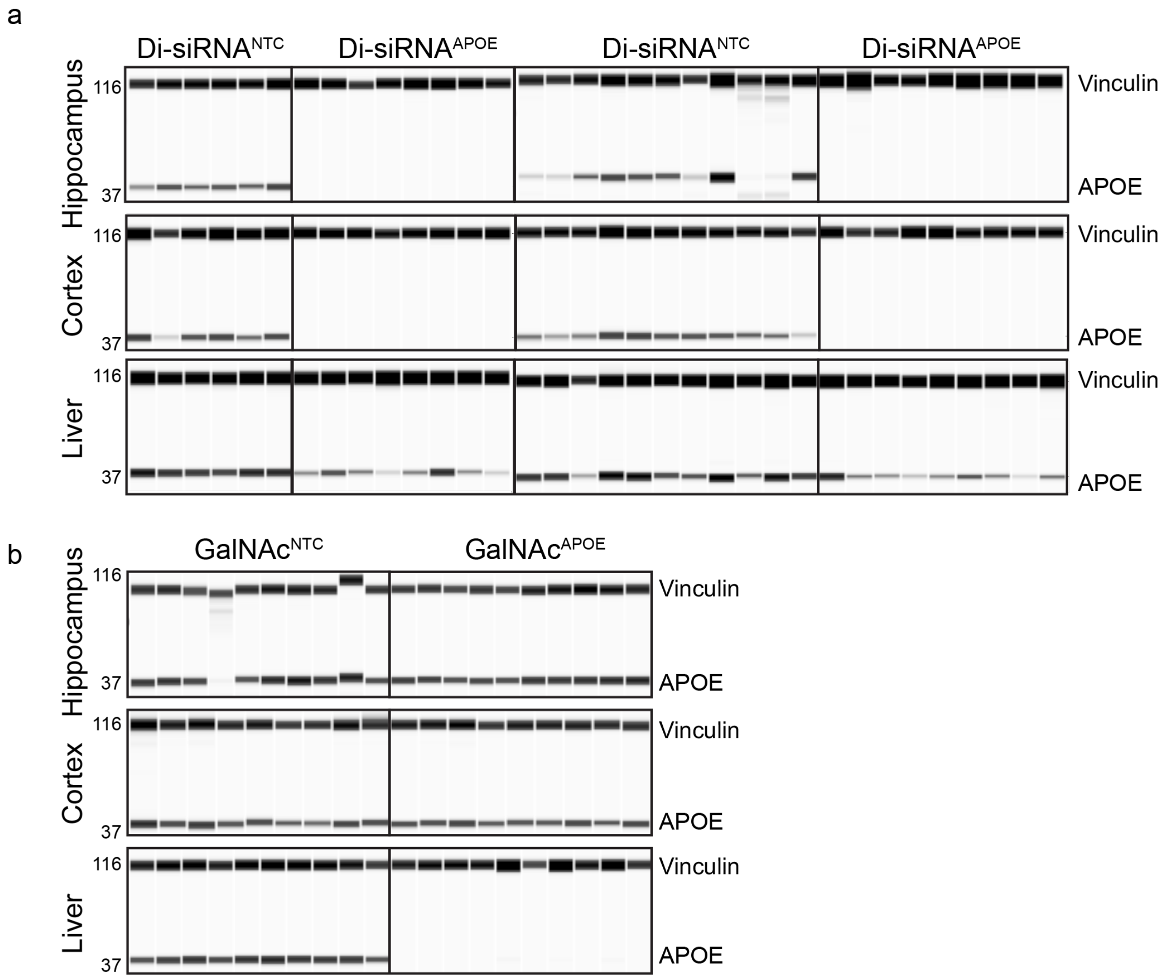


**Extended Figure 2: Liver and CNS *ApoE* protein do not traffic between CNS and Liver in APP/PSEN1 mice. (a)** Raw western blots showing ApoE protein in the hippocampus, cortex, and liver (top to bottom) after administration of di-siRNA$^{NTC}$ or di-siRNA$^{APOE}$. (**b**) Raw western blots showing ApoE protein in the liver but not the hippocampus or cortex after administration of GalNAc$^{NTC}$ or GalNAc$^{APOE}$. ApoE 37 kDa, bottom normalized to loading Vinculin, 116 kDa; top). Protein evaluated with WES for Protein Simple.

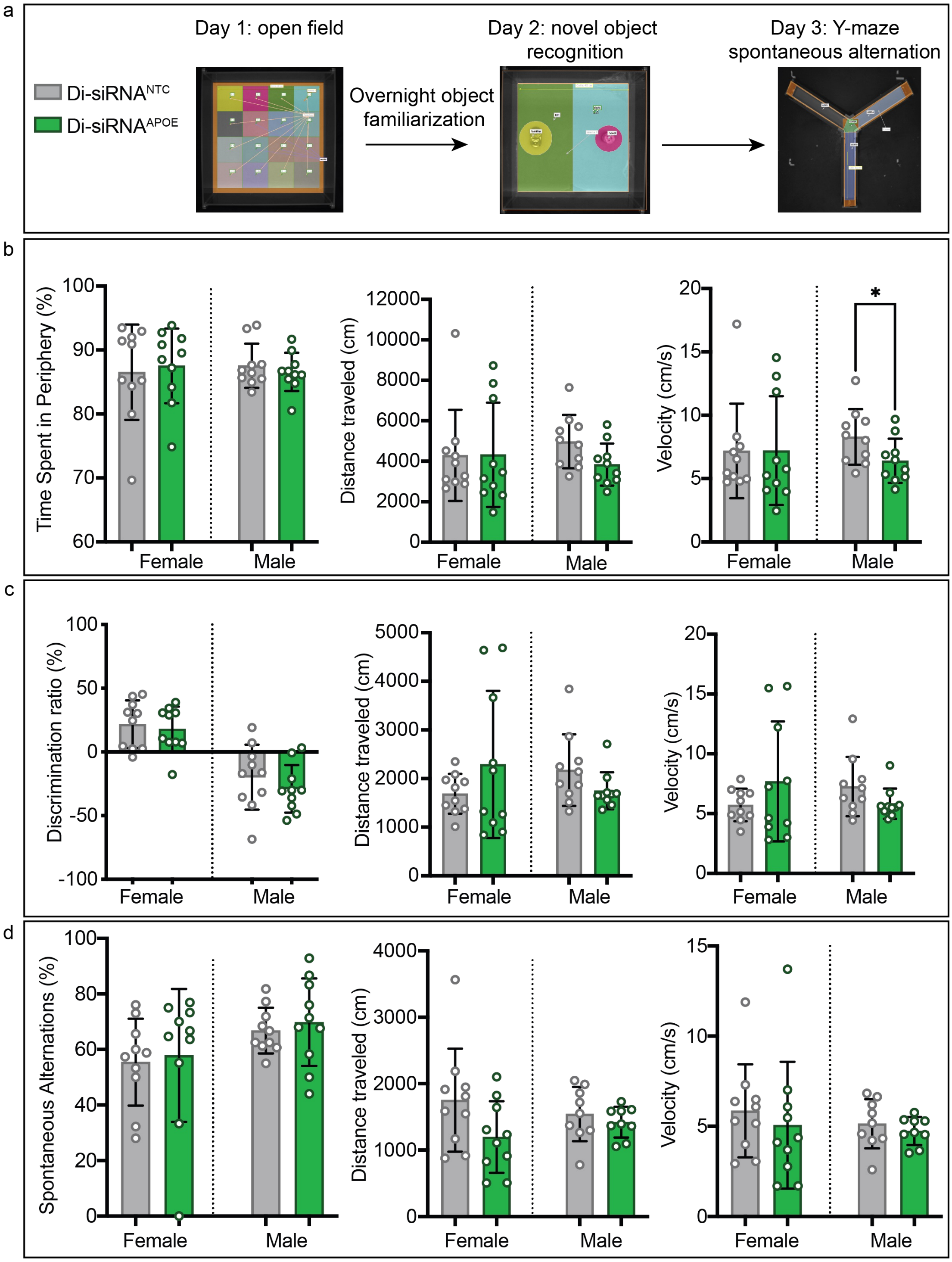


**Extended Data Figure 3: Silencing ApoE with di-siRNA$^{APOE}$ for 2-months does not affect cognitive function in 5xFAD mice. (a)** Study design. Day 1: open field testing. Day 2: Novel Object Recognition. Day 3: y-maze. (**b**) Time spent in periphery (left), distance traveled (middle), and average velocity (right) during open field testing. (**c**) Discrimination ratio (left), distance traveled (middle), and average velocity (right) during novel object recognition. (**d)** Spontaneous alternation (left), distance traveled (middle), and average velocity (right) during open field testing. Experiments performed and analyzed using Noldus EthosVision XT15 software. Statistics performed using GraphPad. T-tests per assay, sex separated. *: p<0.0332; ** p<0.0021; *** p<0.0002; **** p<0.0001. n=9-10 per sex per group. Timepoint: 16-weeks-old, 2 months post injection. Di-siRNA$^{NTC}$ shown in grey, and di-siRNA$^{ApoE}$ shown in green**.**

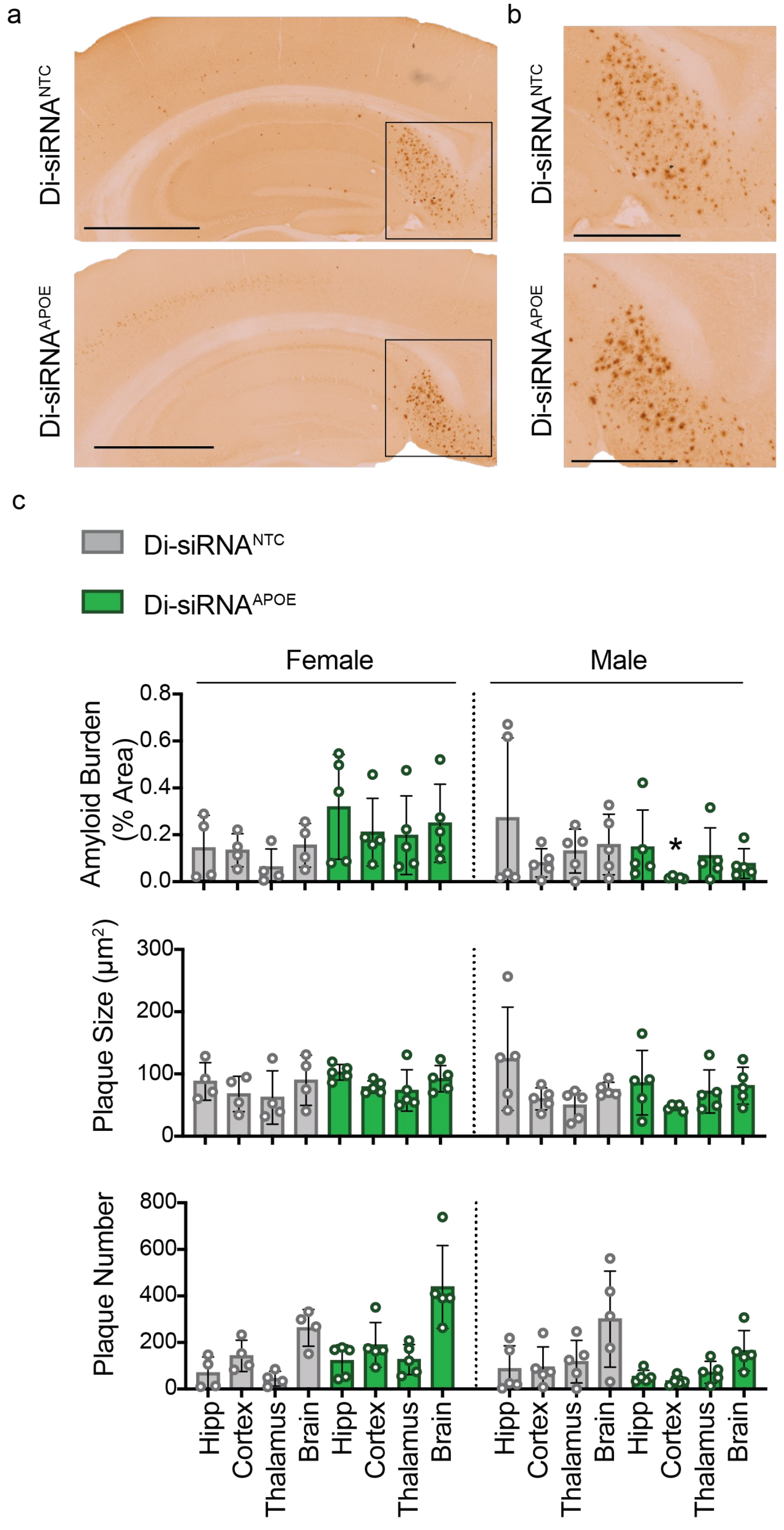


**Extended Data Figure 4: Short-term Silencing of *ApoE* does not reduce Aβ burden.** (**a**) Immunohistochemistry showing amyloid burden (APP 6E10) in control (di-siRNA$^{NTC}$; top) and ApoE silenced (di-siRNA$^{APOE}$; bottom) 5xFAD animals. Scale bar: 1000 µm (**b**) High-resolution images showing onset of plaque deposition in the hippocampus and cortex. Scale bar: 500 µm. (**c**) Quantification of amyloid burden (top), amyloid plaque size (middle), and amyloid plaque number (bottom). Age: 11 weeks; 2 weeks post injection. Female: NTC: n=4/group, APOE: n=5/group; Male: NTC: n=5/group, APOE: n=5/group. Statistics: GraphPad Prism: t-tests per brain region. *: $p<0.0332$; ** $p<0.0021$; *** $p<0.0002$; **** $p<0.0001$. Di-siRNA$^{NTC}$ shown in grey, and di-siRNA$^{APOE}$ shown in green.

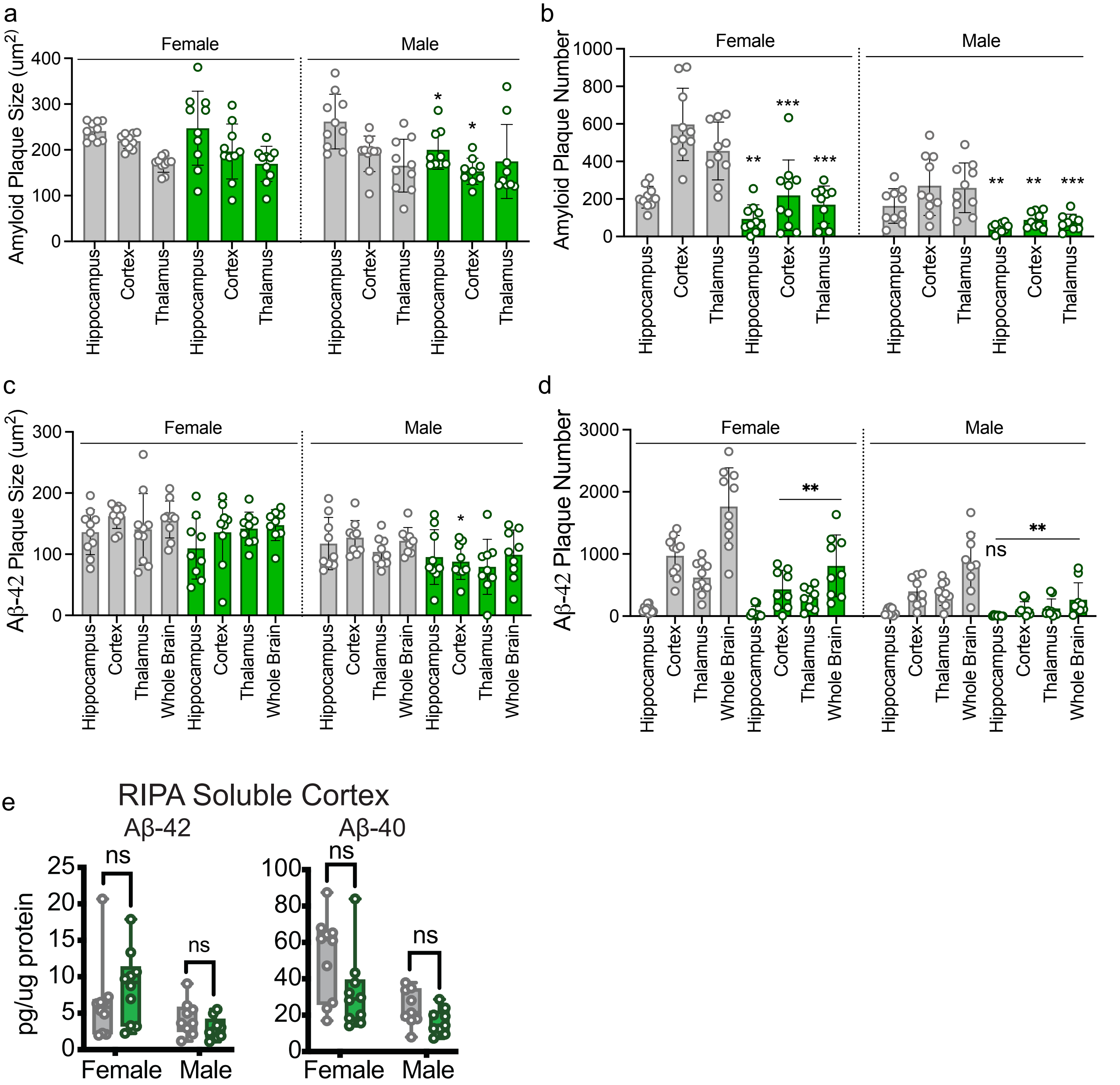


**Extended Data Figure 5: Silencing *ApoE* primarily reduces plaque number not size. (a)** Amyloid plaque size and (**b**) Amyloid plaque number measured using APP6E10 antibody. (**c**) Aβ-42 plaque size and (**d**) Aβ-42 plaque number measured using Aβ-42-specific antibody (D9A3A) in 5xFAD treated and control mice, separated by sex. (**e**) MSD ELIDA quantification of Aβ-42 and Aβ-40 in soluble brain fractions. Statistics: t-test per brain region; sex separated. *: $p<0.0332$; ** $p<0.0021$; *** $p<0.0002$; **** $p<0.0001$. n=9-10 per sex per group. Timepoint: 17 weeks old, 2 months post injection. Di-siRNA$^{NTC}$ shown in grey, and di-siRNA$^{ApoE}$ shown in green.

a

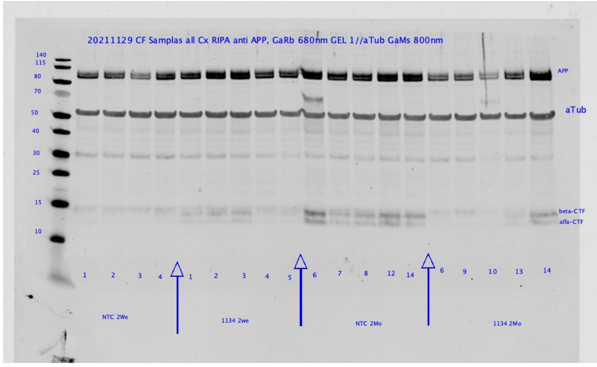


b

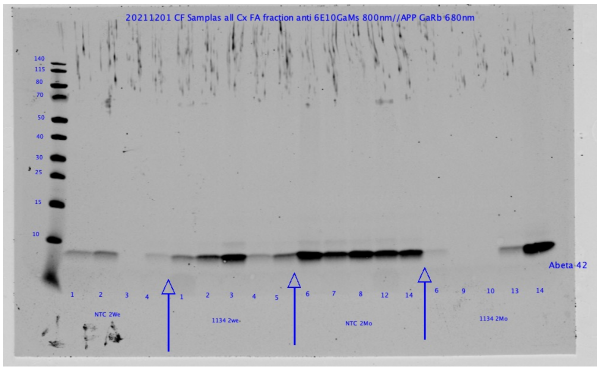


c

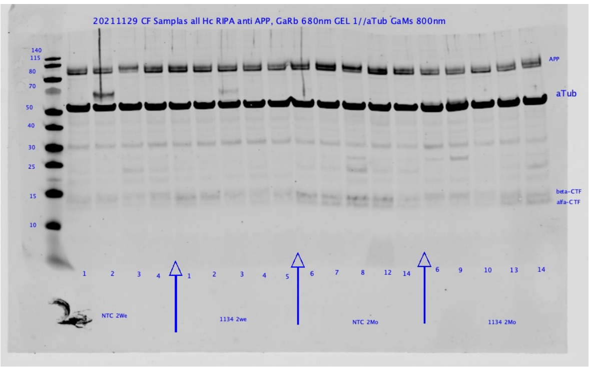


d

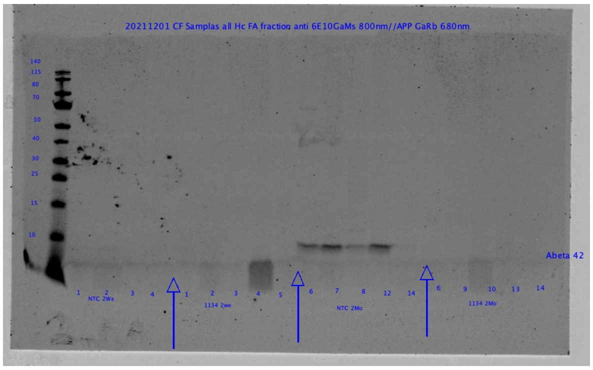


**Extended Data Figure 6: Raw western blot files for analysis of APP, CTFs, and Aβ42 in 5xFAD mice.** (**a**), (**b**) Raw western blots of cortex in a subset of 5xFAD samples. (**c**), (**d**) Raw western blots of hippocampus in a subset of 5xFAD samples. The last lane on all blots is a repeated internal control sample. N=4/5 per group.

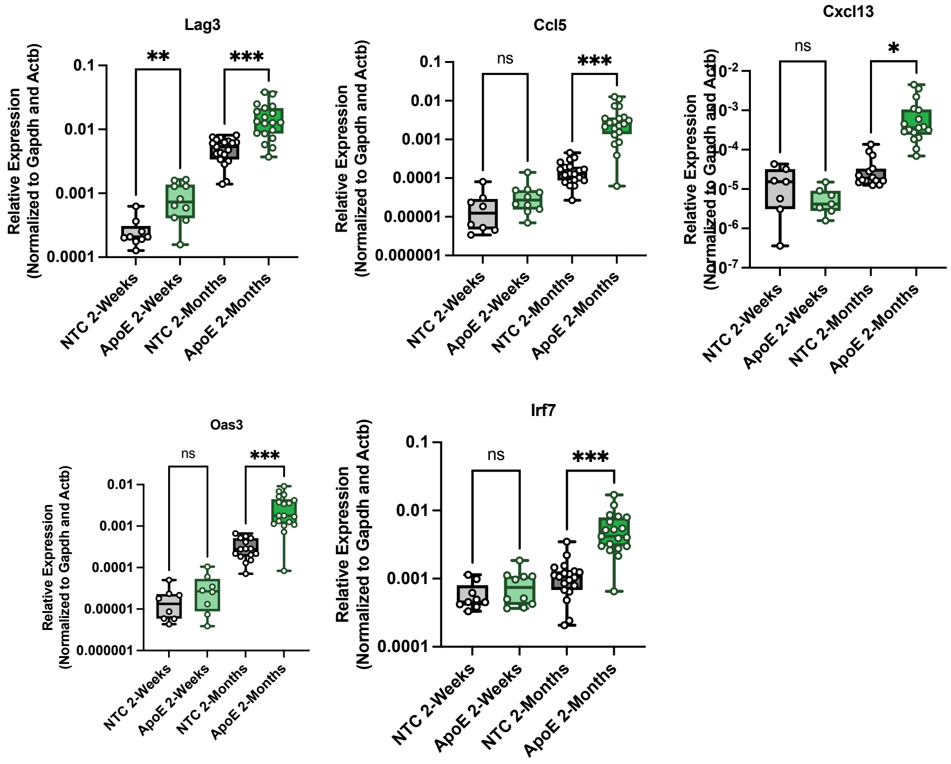


**Extended Data Figure 7: Confirmation of RNA expression using RTqPCR.**
Statistics: One-way ANOVA. *: p<0.0332; ** p<0.0021; *** p<0.0002; **** p<0.0001. n=10-20 per group. Di-siRNA$^{NTC}$ shown in grey, and di-siRNA$^{ApoE}$ shown in green.

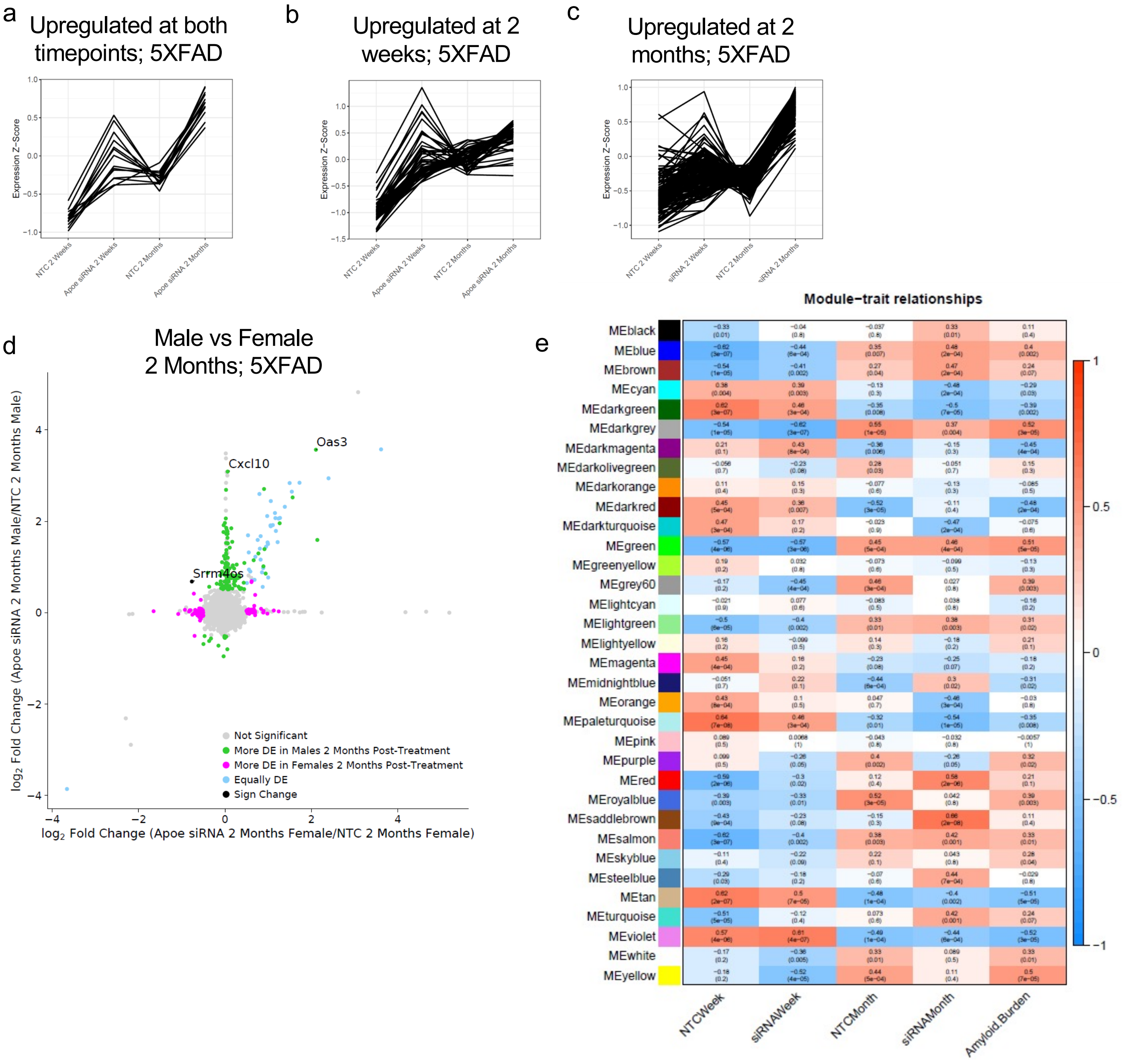


**Extended Data Figure 8: Further analysis of RNA Sequencing: (a-c)** Mean expression z-scores for genes that are upregulated with *ApoE* knockdown at both timepoints (**a**), 2-weeks (**b**) or 2 months (**c**) post-treatment. (**d**) Comparison of differentially expressed genes in male and female 5XFAD mice at 2 months post-treatment. (**e**) Correlation between WGCNA clusters and either treatment or amyloid burden.

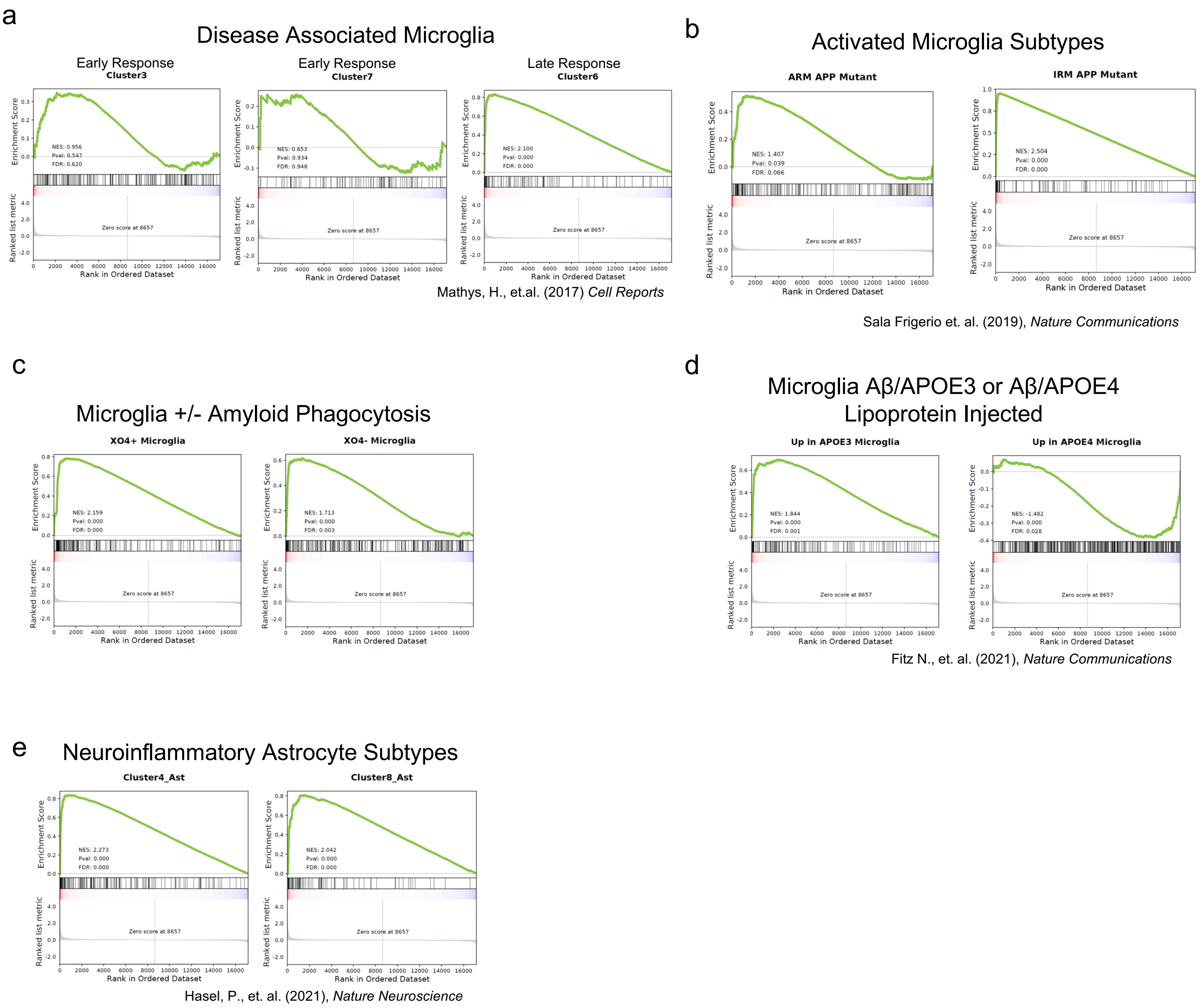


**Extended Data Figure 9: (a)** Comparison of gene overlap between *ApoE* silencing for 2-months in 5XFAD and marker genes of disease associated microglia (Mathys et al., 2017) (**b**), activated and interferon response microglia subtypes (Sala Frigerio et. al. 2019), and (**d**) plaque associated microglia, (**e**) microglial responses after Aβ/APOE3 or Aβ/APOE4 injection (Fitz et al., 2021), and (**f**) marker genes of astrocytes (Hasel et al., 2021).
